# Role of channels in the O_2_ permeability of murine red blood cells III. Mathematical modeling

**DOI:** 10.1101/2025.03.05.639964

**Authors:** Rossana Occhipinti, Pan Zhao, Fraser J. Moss, Walter F. Boron

## Abstract

In this third of three papers, we develop a reaction-diffusion model for O_2_ offloading from a red blood cell (RBC), treated as a sphere with diameter approximating RBC thickness. Stopped-flow (SF) analysis (paper #1) of hemoglobin/oxyhemoglobin (Hb/HbO_2_) absorbance spectra during O_2_ efflux from intact murine RBCs show that membrane-impermeant inhibitor p-chloromercuribenzenesulfonate (pCMBS) reduces the HbO_2_-deoxygenation rate constant (*k*_HbO2_) by ∼61%. SF experiments show that *k*_HbO2_ falls by (1) 9% for aquaporin-1 knockouts (AQP1-KOs), (2) 17% for Rhesus A-glycoprotein knockouts (RhAG-KOs), (3) 30% for double knockouts (dKOs), and (4) ∼78% in dKOs/pCMBS. Here, we simulate HbO_2_ dissociation in the intact RBC (i.e., *k*_HbO2_); HbO_2_, Hb, and O_2_ diffusion through RBC cytosol; transmembrane O_2_ diffusion; and O_2_ diffusion through extracellular unconvected fluid (EUF) to bulk extracellular fluid. Informed by automated-hematology data (paper #1) and imaging–flow-cytometry data (paper #2), simulations predict that observed *k*_HbO2_ decreases cannot reflect changes only in RBC size/shape or [Hb/HbO_2_]. Instead, membrane O_2_ permeability (*P*_M,O2_) must fall by (1) 22% to account for AQP1-KO data, (2) 36% for RhAG-KOs, (3) 55% for dKOs, and (4) 91% for dKOs/pCMBS. Exploring predicted *k*_HbO2_ sensitivities to eight key parameters (e.g., [Hb/HbO_2_], diffusion constants, *k*_HbO2→Hb+O2_, thickness_EUF_, diameter_Sphere_) shows that no reasonable changes explain the *k*_HbO2_ data. We introduce a linear-combination approach to accommodate for the presence of poikilocytes. Finally, contrary to common beliefs, the model predicts that, in the absence of inhibitors, the RBC membrane represents >30% of total diffusive “resistance” to O_2_ offloading, even for a WT mouse.

**Key Points:**

- In this third of three papers, we develop a novel reaction-diffusion model for O_2_ offloading from a red blood cell (RBC), treated as a sphere with diameter approximating RBC thickness.
- Using physical constants and parameter values from the literature and papers #1 and #2, we generate simulations that reproduce observed hemoglobin desaturation (rate constant, *k*_HbO2_) of intact RBCs from wild-type mice.
- To simulate *k*_HbO2_ decreases (9%, 17%, 30%) observed in RBCs from knockouts (KOs: aquaporin-1, Rhesus A-glycoprotein, both) we must decrease membrane O_2_ permeability (*P*_M,O2_) by far greater percentages (22%, 36%, 55%).
- Analyses of simulated-*k*_HbO2_ sensitivity to kinetic and geometric parameters suggest that reasonable parameter-value changes cannot explain experimentally observed *k*_HbO2_ decreases in RBCs from KOs. Thus, experimentally observed *k*_HbO2_ decreases must reflect decreases in *P*_M,O2_.
- To accommodate for poikilocytes, we develop a linear-combination approach (poikilocytes + biconcave disks, BCDs) to extract the *k*_HbO2_ and *P*_M,O2_ of just BCDs.

## Introduction

Red blood cells (RBCs) are uniquely suited to perform the vital task of taking up oxygen (O_2_) in the pulmonary capillaries, carrying this O_2_ to the systemic capillaries, and there offloading the O_2_ for consumption in the peripheral tissues. This journey of O_2_ within the vertebrate body— and the oppositely directed journey of carbon dioxide (CO_2_)—involves a series of convective and diffusive steps within and across the various components of the respiratory system.

Integral to the above sequence of events is an important series of membrane-permeation steps, both for O_2_ delivery and CO_2_ removal. Among the membranes that O_2_ and CO_2_ must traverse is the plasma membrane of RBCs. The mechanisms by which these gases move across cell membranes, including the RBC membranes, have been the subject of numerous investigations (Graham, 1829, 1833; Mitchell, 1830, 1833; Fick, 1855; Overton, 1895; Wroblewski, 1879; Hartridge & Roughton, 1927; Forster, 1964; Waisbren *et al*., 1994; Nakhoul *et al*., 1998; Cooper & Boron, 1998) and reviews (Stannett, 1978; Geers & Gros, 2000; Boron, 2010; Boron *et al*., 2011; Endeward *et al*., 2014; Cooper *et al*., 2015; Michenkova *et al*., 2021). Prior to the 1990s, two major schools of thought emerged. One, influenced by Krogh’s cylinder model of O_2_ diffusion from capillary to tissue, implicitly assumes that the RBC membrane offers no resistance to O_2_ diffusion (Krogh, 1919; Kreuzer, 1982)—that is, the membrane behaves as a layer of water of equivalent thickness. The other school of thought, influenced by Overton’s work on the lipoid nature of plasma membranes, is that the permeability of a membrane to a solute like O_2_ depends only on the solubility of the solute in the lipid phase of the membrane (Overton, 1895, 1899).

The present contribution is the third in a series of three interconnected papers^1^ that investigate the hypothesis that two membrane proteins that are abundant in the murine RBC membrane, aquaporin-1 (AQP1) and the Rh complex (Rh_Cx_)—comprising Rhesus blood group-associated A-glycoprotein (RhAG) and + Rhesus blood group D antigen (RhD)—represent a large fraction of the membrane permeability to O_2_ (*P*_M,O2_).

In the first paper (<u>paper #1</u>; Zhao *et al*., 2025), we employ stopped-flow (SF) analysis of hemoglobin/oxyhemoglobin (Hb/HbO_2_) absorbance spectra to measure the rate constant of Hb deoxygenation (*k*_HbO2_) in RBCs from mice genetically deficient in AQP1 (*Aqp1−/−*), RhAG (*Rhag−/−*), or both (*Aqp1−/−Rhag−/−*). In both wild-type (WT) mice and double knockouts (dKO), we also examine the effects of the inhibitors p-chloromercuribenzenesulfonate (pCMBS) and 4,4’-diisothiocyanatostilbene-2,2’-disulfonate (DIDS). Because *k*_HbO2_ depends on factors other than *P*_M,O2_, we also assess in paper #1 the rate constant of the dissociation HbO_2_ → Hb + O_2_ in RBC hemolysates (*k*_HbO2→Hb+O2_), and automated-hematological parameters, including mean corpuscular volume (MCV), mean corpuscular hemoglobin (MCH), and mean corpuscular hemoglobin concentration (MCHC).

We find that—compared to WT mice—*k*_HbO2_ decreases by 9% in RBCs from AQP1-KOs, by 17% in RhAG-KOs, and by 30% in dKO. This work thus points towards roles for AQP1 and RhAG in enhancing O_2_ permeation through RBC membranes.

In the second paper (i.e. <u>paper #2</u>; Moss *et al*., 2025),, we examine RBC morphology at a microscopic scale, and confirm that the RBCs from the KO and dKO mice are predominantly biconcave disks (BCDs). We also use imaging flow cytometry (IFC) to assess RBC major diameter (Ø_Major_). Together with MCV data from paper #1, Ø_Major_ allows us to estimate RBC thickness—a critical piece of information for the simulations in this third (the present) paper in the series. The IFC data also shows that a small fraction of RBCs, more so in drug-treated cells, are poikilocytes, that is, non-biconcave disks (nBCDs). This is also critical information for the present paper because it sets the stage for a novel correction methodology that we apply here. Finally, paper #2 includes a proteomic study that shows that the KO of one or both channels does not affect the apparent abundance of any of the 100 proteins with the greatest inferred abundance.

The purpose of this third (the present) paper in the series is to develop a reaction-diffusion model of O_2_ offloading from RBCs, and to use this model—based only on first principles and informed by MCV, MCH, MCHC, *k*_HbO2→Hb+O2_, and Ø_Major_ data from the first two papers—to simulate the SF data from the paper #1. We find that the simulations agree reasonably well with the time course of deoxygenation for RBCs from WT mice (measured as *k*_HbO2_ values) in paper #1. Note that our approach is not merely to adjust a variety of fitting parameters in an effort to reproduce the observed *k*_HbO2_. Except for sensitivity analyses—when we examine how sensitive the simulations are to variations in specific parameters—the only parameter that we vary is *P*_M,O2_, which we do to predict the *P*_M,O2_ values that reproduce the experimentally determined time courses. Our graphical user interface (Huffman *et al*., 2025) expands the power of the model, especially for non-experts.

Taken together, the three papers argue strongly against each of two previously dominant— and quite opposite—schools of thought (addressed in the Discussion). One holds that RBC membranes offer no resistance whatsoever to O_2_ diffusion. The other, posits that they do offer resistance, but that this resistance depends solely on the solubility of O_2_ in RBC membrane lipids. Instead, our data are consistent with the notion that RBC membranes from WT mice offer ∼150-fold more resistance to O_2_ diffusion than an equivalently thick layer of water, and that channels—AQP1, RhCx, and an unidentified channel(s)—account for >90% of *P*_M,O2_ under the conditions of our experiments^2^.

## Methods

Figure 1 shows the workflow for the three-paper project. The present paper focuses on steps #10, #12 and #12′, #13, #14 and #14′, #16, #17 and #17′, #18 through #22.

**Figure 1.**
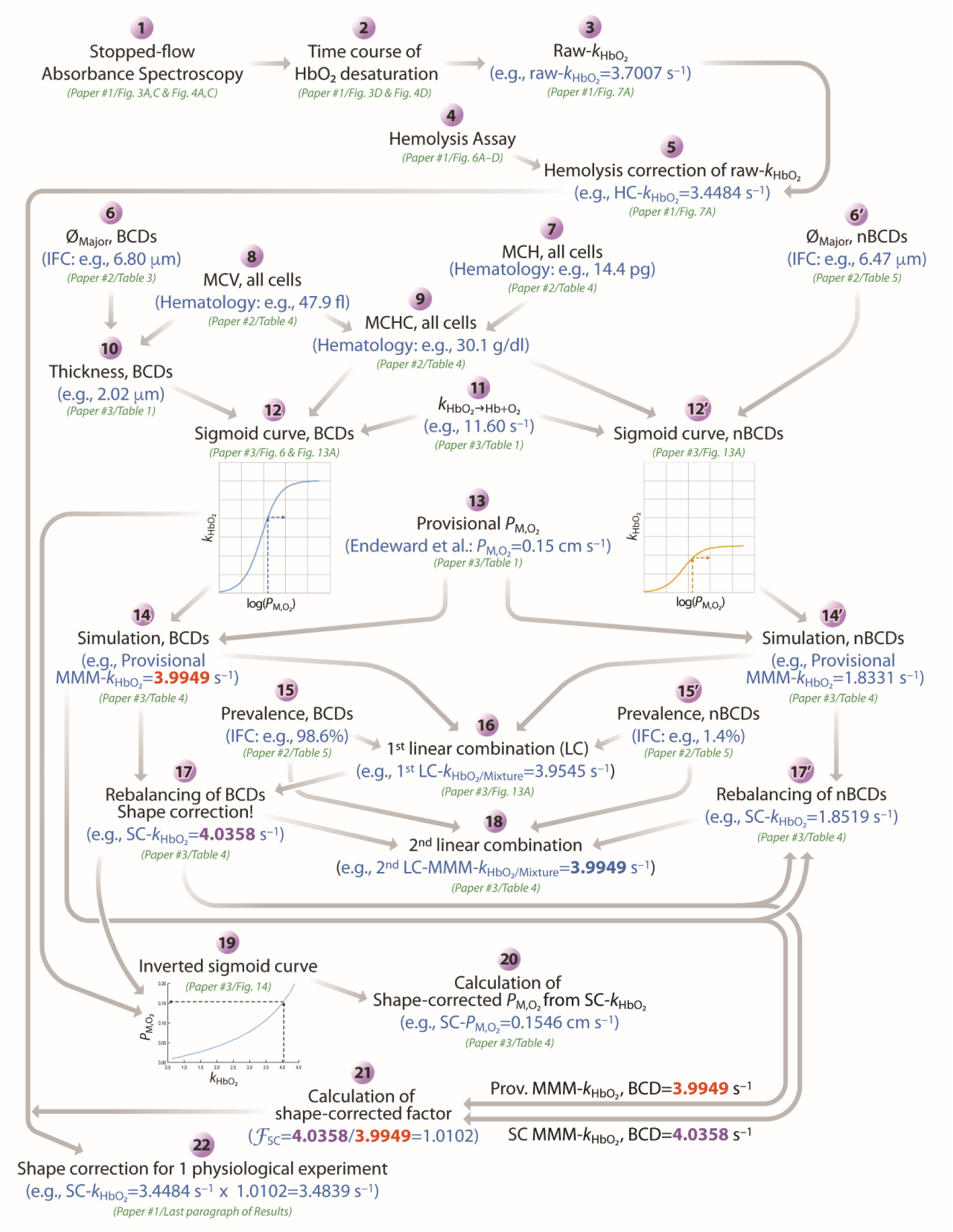
Workflow The numerals 1 through 22 indicate the steps summarized in <u>paper #1</u>—and presented in detail, as appropriate, in the present paper (<u>paper #3</u>), <u>paper #1</u>, or <u>paper #2</u>—to arrive, first, at a hemolysis-corrected (HC) *k*_HbO2_ and, ultimately, at a shape-corrected (SC) *k*_HbO2_. At each step, we provide example values, if possible, and example figure/table numbers referenced to <u>paper #1</u>, <u>paper #2</u>, or the present paper. We repeat workflow steps 1–5 for each mouse, ultimately arriving at an HC-*k*_HbO2_ value for each. Step 6 applies to biconcave disks (BCDs), whereas step 6′ applies to non-biconcave disks (nBCDs). The same is true for steps 12 vs. 12′, 14 vs. 14′, 15 vs. 15′, and for 17 vs. 17′. *F*_SC_, shape-correction factor; Hb, hemoglobin; IFC, imaging flow cytometry; MCH, mean corpuscular hemoglobin; *k*_HbO2_, rate constant for deoxygenation of HbO_2_ within intact RBCs; *k*_HbO2→Hb+O2_, rate constant for deoxygenation of HbO_2_ in free solution; LC, linear combination; MCH, mean corpuscular hemoglobin; MCHC, mean corpuscular hemoglobin concentration; MCV, mean corpuscular volume; MMM-k_HbO2_, macroscopic mathematically modeled rate constant for deoxygenation of intracellular HbO_2_; *P*_M,O2_, membrane permeability to O_2_; Prov., provisional; Ø_Major_, major diameter (of BCD or nBCD); SC-k_HbO2_, shape-corrected rate constant for deoxygenation of intracellular HbO_2_.

### Mathematical modeling and simulations

#### Model formulation

To examine the possibility that changes in RBC parameters—decreases in *k*_HbO2→Hb+O2_, increases in MCHC (i.e., [Hb]_i_ + [HbO_2_]_i_ = [Hb_Total_]_i_)^3^, decreases in diffusion constants (i.e., intra-/extracellular *D* for O_2_; intracellular *D* for HbO_2_, Hb), increases in the thickness of the unconvected-fluid layer that surrounds the cell, or increases in intracellular diffusion distances—can explain our observed decreases in *k*_HbO2_, we developed a distributed (i.e., space-dependent) and dynamic (i.e., time-dependent) reaction-diffusion mathematical model of an RBC—simplified as a perfectly symmetric sphere with diameter equal (Ø_Sphere_) to the thickness of the RBC. Previous modelers have used a sphere in their simulations of an RBC (Vandegriff & Olson, 1984*a*; Williams & Kutchai, 1986; Deonikar & Kavdia, 2013; Deonikar *et al*., 2014), matching the diameter of the sphere to Ø_Major_. We avoid equating our Ø_Sphere_ with the major diameter of the BCD because this would generate unrealistically large diffusion distances and thus low values of *k*_HbO2_ and *P*_M,O2_. As described below (see <u>below</u>^4^), our geometrical approach is also unique in that we chose Ø_Sphere_ to match the minor diameter of a torus that has the same volume as the RBC. In our model (see Figure 2A), the spherical RBC is enveloped by a layer of extracellular unconvected fluid (EUF) that is in turn enveloped by the bulk extracellular fluid (bECF). The model accounts for diffusion of O_2_, Hb, and HbO_2_ within the intracellular fluid (ICF). Only O_2_ can move—via diffusion—between the ICF and EUF (i.e., across the plasma membrane, M), within the EUF (which lacks Hb and HbO_2_), and between the EUF and bECF (which also lacks Hb and HbO_2_). Reactions among O_2_, Hb, and HbO_2_ occur only in the ICF.

**Figure 2.**
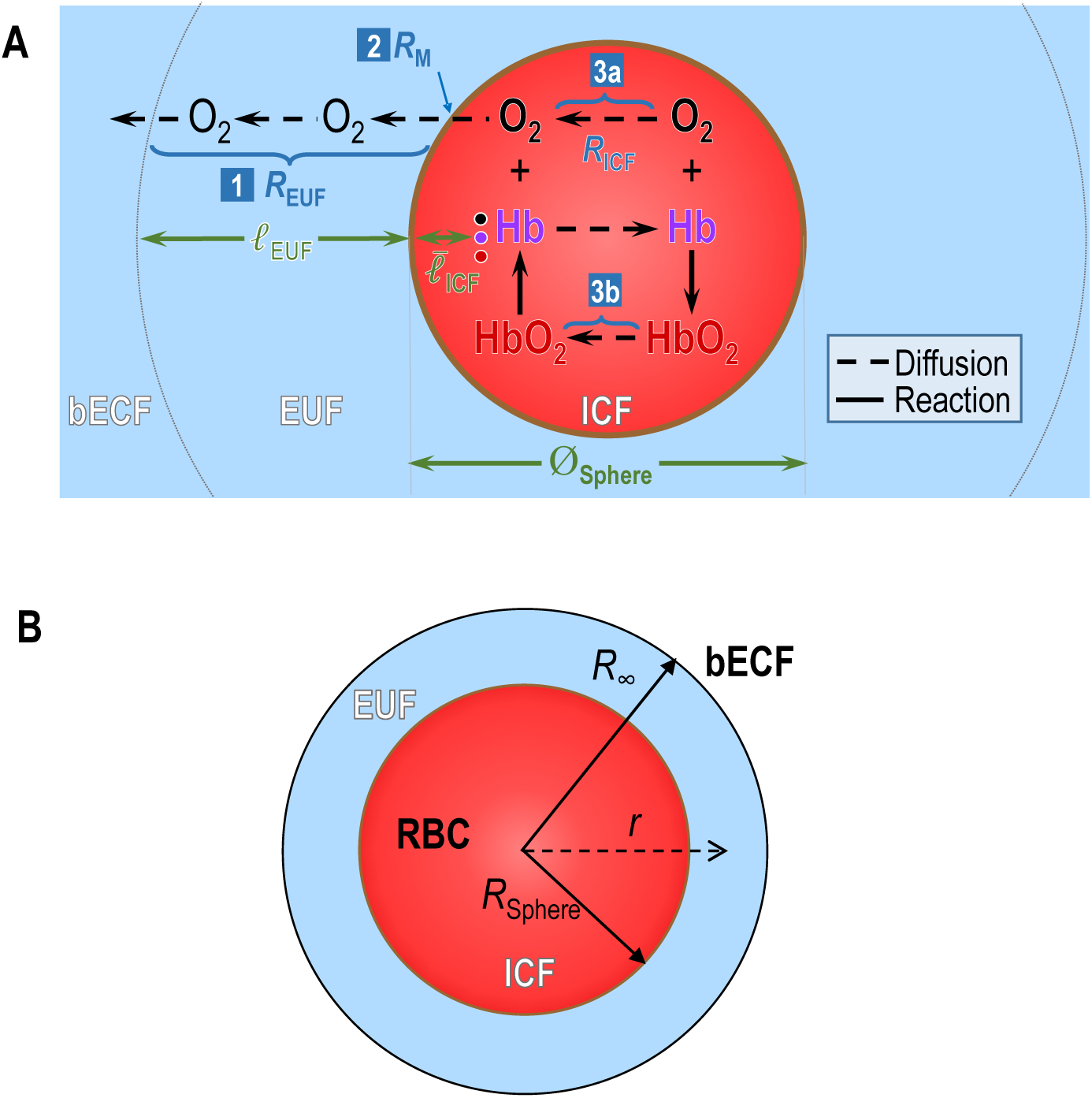
Mathematical model of O_2_ efflux from an RBC *A,* Major components of the mathematical model of spherical RBC cell, the diameter of which matches the computed thickness of an average RBC from a WT or KO mouse. See text for details on the model. *B,* The computational domain. For panel ‘A’, the abbreviations are: bECF, bulk extracellular fluid; EUF, extracellular unconvected fluid; ICF, intracellular fluid; M, plasma membrane; Ø_Sphere_, diameter of sphere (matches thickness of RBC); ℓ_EUF_, thickness of EUF; 3 values of are average distance from M to the following 3 molecules in ICF: (a) O_2_ (black dot and lettering), (b) Hb (violet), or HbO_2_ (red) in ICF; [1] *R*_EUF_, effective resistance offered by EUF to O_2_ diffusion from M to bECF; [2] *R*_M_, effective resistance offered by M to O_2_ diffusion; [3] *R*_ICF_, effective resistance offered by ICF to combined diffusion of O_2_ and HbO_2_. 1/*R*_ICF_ = 1/*R*_ICFa_ + 1/*R*_ICFb_, where *R*_ICFa_ is effective resistance offered by ICF to O_2_ diffusion over the distance ^−^_ICF_, from the position of the average O_2_ to M; *R*_ICFb_ is effective resistance offered by ICF to HbO_2_ diffusion over the distance ^−^_ICF_, from the position of the average HbO_2_ to M. For panel ‘B’, the abbreviations are: *R*_∞_, maximal radial distance from the center of the sphere (*R*_∞_ = *R*_sphere_ + ℓ_EUF_); *r*, radial distance from the center of the cell.

Assuming spherical-radial symmetry, then for each solute *S* (i.e., O_2_, Hb, and HbO_2_), the concentration (*C*_S_) changes in time (*t*) and space according to the reaction-diffusion equation

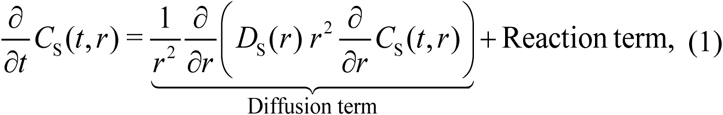

where *r* (without a subscript^5^) is the radial distance from the center of the RBC, and *D_S_* is the diffusion coefficient of solute *S*. Figure 2B shows the computational domain.

#### Kinetics of the oxygen-hemoglobin reaction

We model the binding of O_2_ to Hb (and dissociation of O_2_ from HbO_2_)—that is, the reaction term in Equation (1)—as a simple, one-step reaction in which we use a version of the variable-rate-coefficient (VRC) approach proposed by Moll (Moll, 1968). Thus, we assume that each hemoglobin tetramer (Hb_T_) is replaced by four independent hemoglobin monomers (4 × Hb_M_), and that each hemoglobin monomer can combine with an O_2_ via the one-step reaction

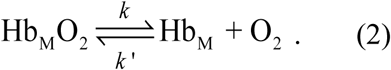

Here, the dissociation rate constant *k* has the same meaning as *k*_HbO2→Hb+O2_ in <u>paper #1</u>^6^, namely, the rate constant for deoxygenation of HbO_2_ in an RBC hemolysate; we will use the two notations interchangeably. The rate constant *k*′ for the association reaction is equivalent to *k*_Hb+O2→HbO2_. Rather than assuming that both *k* and *k′* are constant—an assumption that would lead to a non-realistic hyperbolic hemoglobin-oxygen saturation curve—we follow a version of Moll’s VRC approach, in which we assume that only one of the two reaction rates is constant, while the other one becomes a function of the partial pressure of O_2_ (P_O2_) so that the mathematical representation of the hemoglobin-oxygen saturation curve corresponds to the measured sigmoidal curve. Yap and Hellums (1987) showed that the VRC model is appropriate in the physiological range, and avoids the use of the more complex Adair model. In his VRC approach, Moll assumed that *k′* is constant and *k* is a function of P_O2_. However, Clark *et al*. (1985) pointed out that the function used by Moll to describe *k* as a function of P_O2_ approaches infinity as P_O2_ approaches zero. These last authors overcame this problem by assuming that *k* is constant and *k′* is a function of P_O2_, as described by

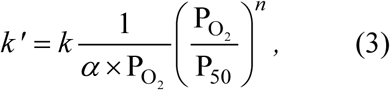

where *α* is the solubility coefficient for O_2_ in water, P_50_ is P_O2_ when [Hb] = [HbO_2_], and *n* is the Hill coefficient. With this choice and *n* > 1, *k*′ approaches zero as P_O2_ approaches zero. Thus, in our model, we follow Clark’s variation of the VRC approach, which we implement with the slight modification of substituting [O_2_] for P_O2_:

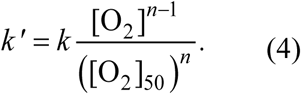

#### Calculation of RBC thickness based on the geometry of a torus

For each mouse strain, we calculate the RBC thickness as follows (Figure 3). We start from the mean major diameter of all RBC-related cell types^7^ (e.g., Ø_Major_ = 6.80 μm for WT cells; see <u>paper #2</u>^8^)—a measured value— and assume that the RBC is a torus, the outer diameter (*OD*_Torus_) of which equals the aforementioned Ø_Major_ (i.e., 6.80 μm in our example). Next, we use the mean corpuscular volume (e.g., MCV = 47.9 pl for WT cells; see <u>paper #2</u>^9^)—also a measured value, which we assume to be equal to the volume of the torus (*V*_Torus_)—to calculate the minor radius (*r*_Torus_) of the torus (equivalent to half the thickness of the RBC). Thus, *r*_Torus_ is a root of the polynomial:

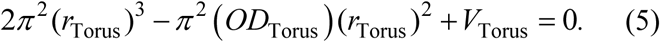

**Figure 3.**
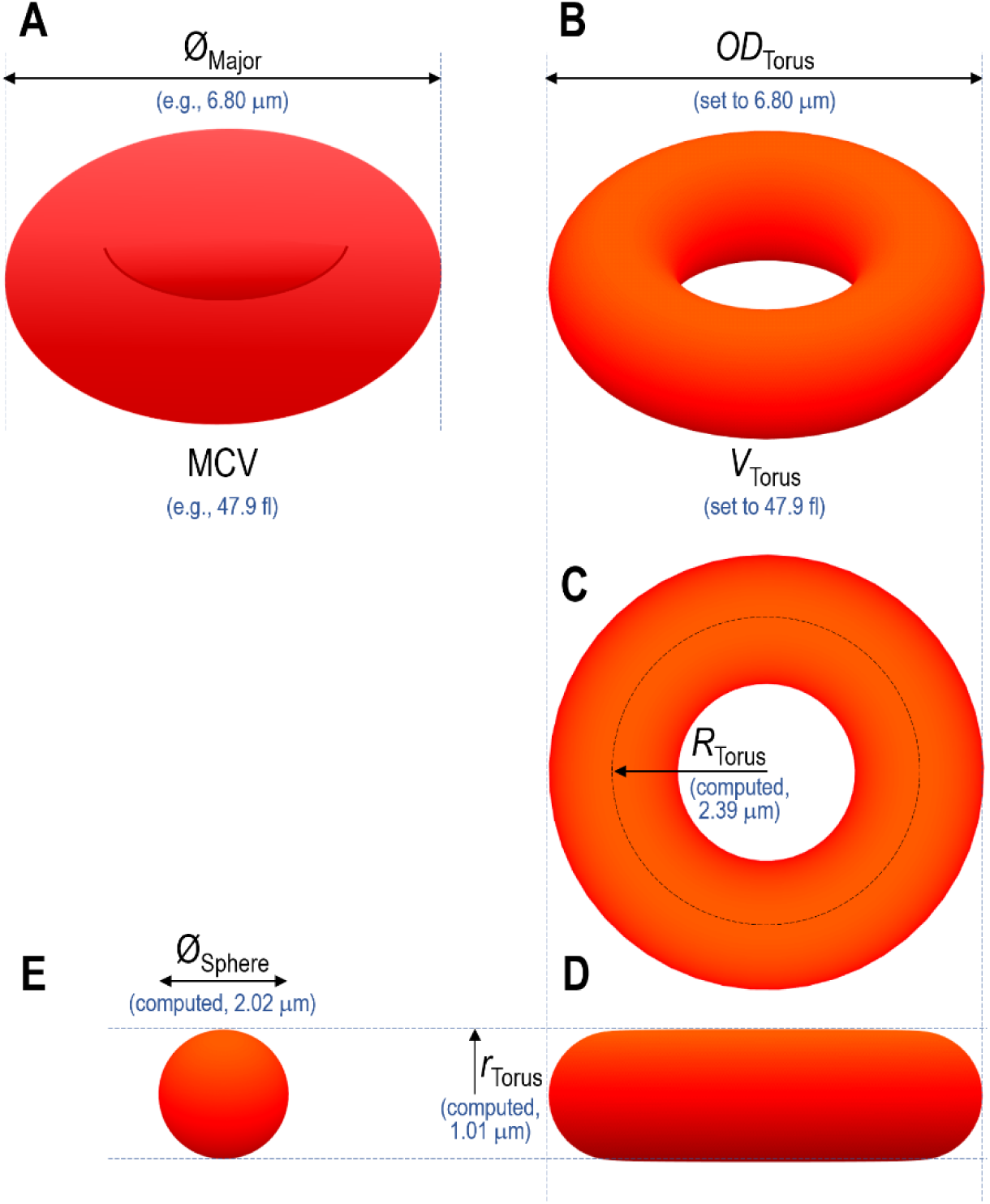
Conversion of biconcave disk geometry into torus geometry *A,* Biconcave disk, tilted forward. *B,* Torus, tilted forward. *C,* Torus, top view. *D,* Torus, side view. *E,* Equivalent sphere. Ø_Major_, major diameter of biconcave disk; Ø_Sphere_, diameter of equivalent sphere; MCV, mean corpuscular volume; *OD*_Torus_, outer diameter of torus; *r*_Torus_, minor radius of torus; *R*_Torus_, major radius of torus; *V*_Torus_, volume of torus (made equal to MCV).

The calculated value for the WT cells is *r*_Torus_ = 1.01 μm. In Table 2, we provide—for each of the 4 mouse strains—the computed values for *r*_Torus_, the major radius of the torus (*R*_Torus_), and the diameter of the equivalent sphere, where Ø_Sphere_ = 2*r*_Torus_ (i.e., RBC thickness). The equivalent sphere diameters serve as inputs to our mathematical model.

#### Calculation of Hb content

For each mouse strain, we calculate the total concentration (both O_2_-bound and free, in mM) of Hb tetramers (T)—that is, ([Hb_T,Total_])^10^ —as follows. Based on the hematology data^11^ reported in <u>paper #2</u>^9^—specifically, the MCH (in pg) and MCV (in fl)—and using a value of 64316 g/mol for the molecular weight of mouse Hb, we calculate the MCHC (in mM). For WT mice, the concentration of total Hb tetramers was ∼4.7 mM. However, because our mathematical model deals with monomers, we calculate the concentration of total of Hb monomers ([Hb_M,Total_])^12^ using 4 × MCHC. In the case of RBCs from WT mice, this figure is <u>18.73 mM</u>. Table 2 lists this value, as well as the comparable values for the other three genotypes. These values inform our mathematical model.

#### Computational model

In order to solve the system of the three reaction-diffusion equations (1)—one for O_2_, one for Hb monomers, and one for HbO_2_ monomers—we set the following boundary conditions.

At the center of the RBC, we posit the no-flux Neumann boundary condition:

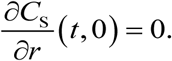

At the interface between the ICF and the EUF (i.e., at the M), we establish continuity of Fick’s diffusive flux:

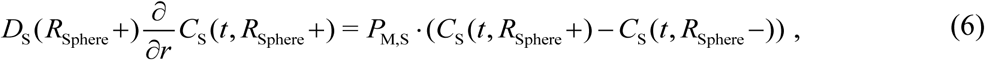

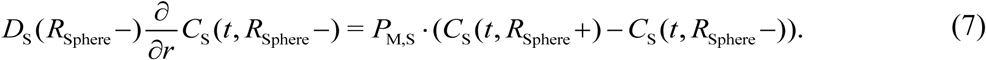

Here, *R*_Sphere_ is the radius of the sphere, *C*_S_(*t*, *R*_Sphere_–) is the time-dependent concentration of solute *S* in the aqueous phase adjacent to the intracellular (−) side of the M; *C_S_*(*t*, *R*_Sphere_+), the comparable value on the extracellular (+) side; and *P*_M*,S*_ is the true permeability^13^ of the membrane to solute *S*.

Because O_2_ is the only solute that can diffuse across the M, *P*_M*,S*_ ≠ 0 only for *S* = O_2_.^14^

In the bECF, we assign Dirichlet boundary conditions and assume that the concentration of O_2_ is constant and equal to zero. This assumption is the mathematical equivalent of the annihilation of O_2_ by sodium dithionite in our SF reaction cell (see <u>paper #1</u>^15^).

Finally, we set the initial conditions by assuming that, at the beginning of the experiment (i.e., our simulation), no O_2_ is present in the EUF, and the free [O_2_] everywhere inside the RBC is that dictated by Henry’s law for a gas mixture containing 21% O_2_. The initial concentrations of O_2_, Hb, and HbO_2_ in the ICF are those reported in Table 1 for RBCs from WT mice, and in Table 2, for RBCs from mice of each of four genotypes.

**Table 1.**
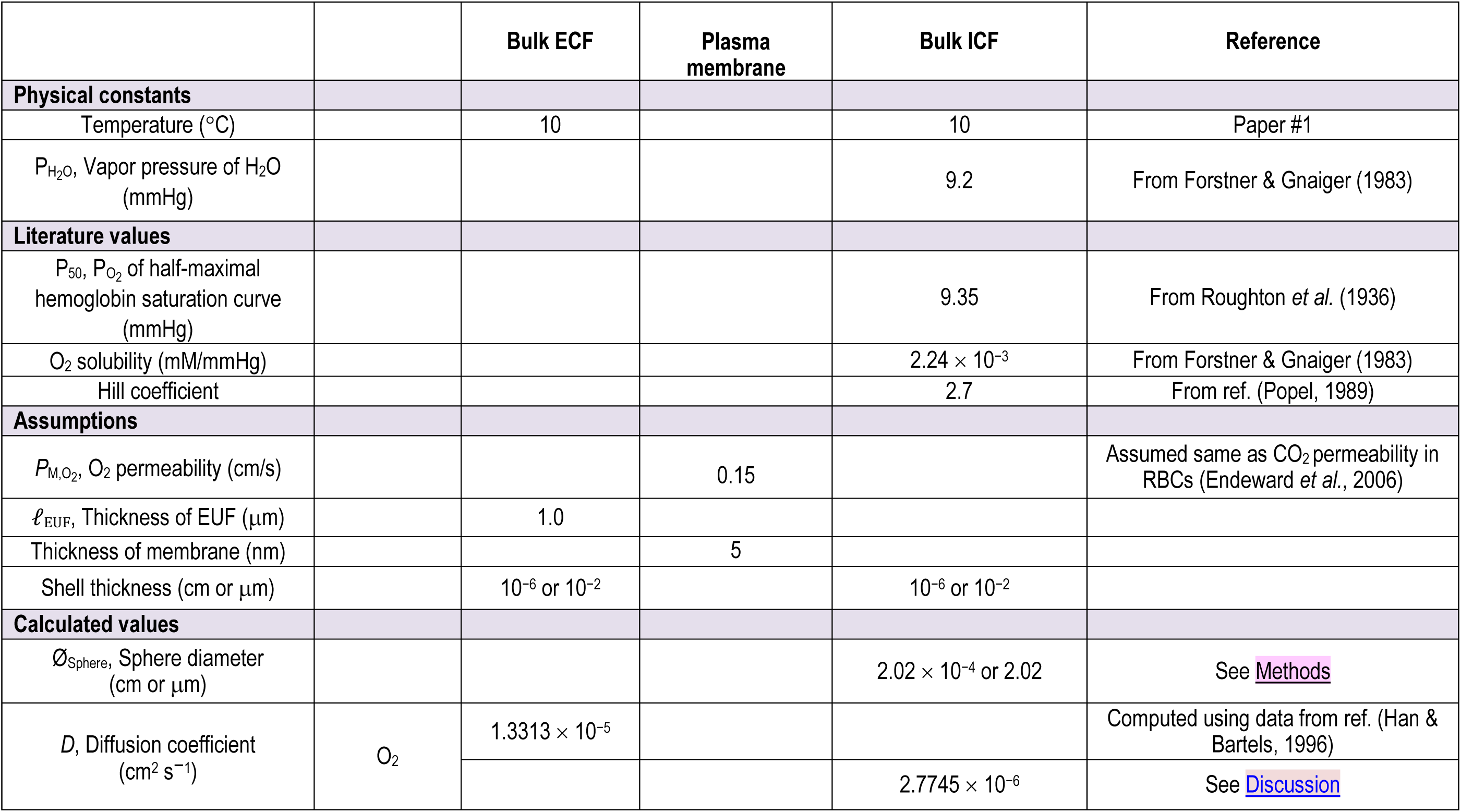
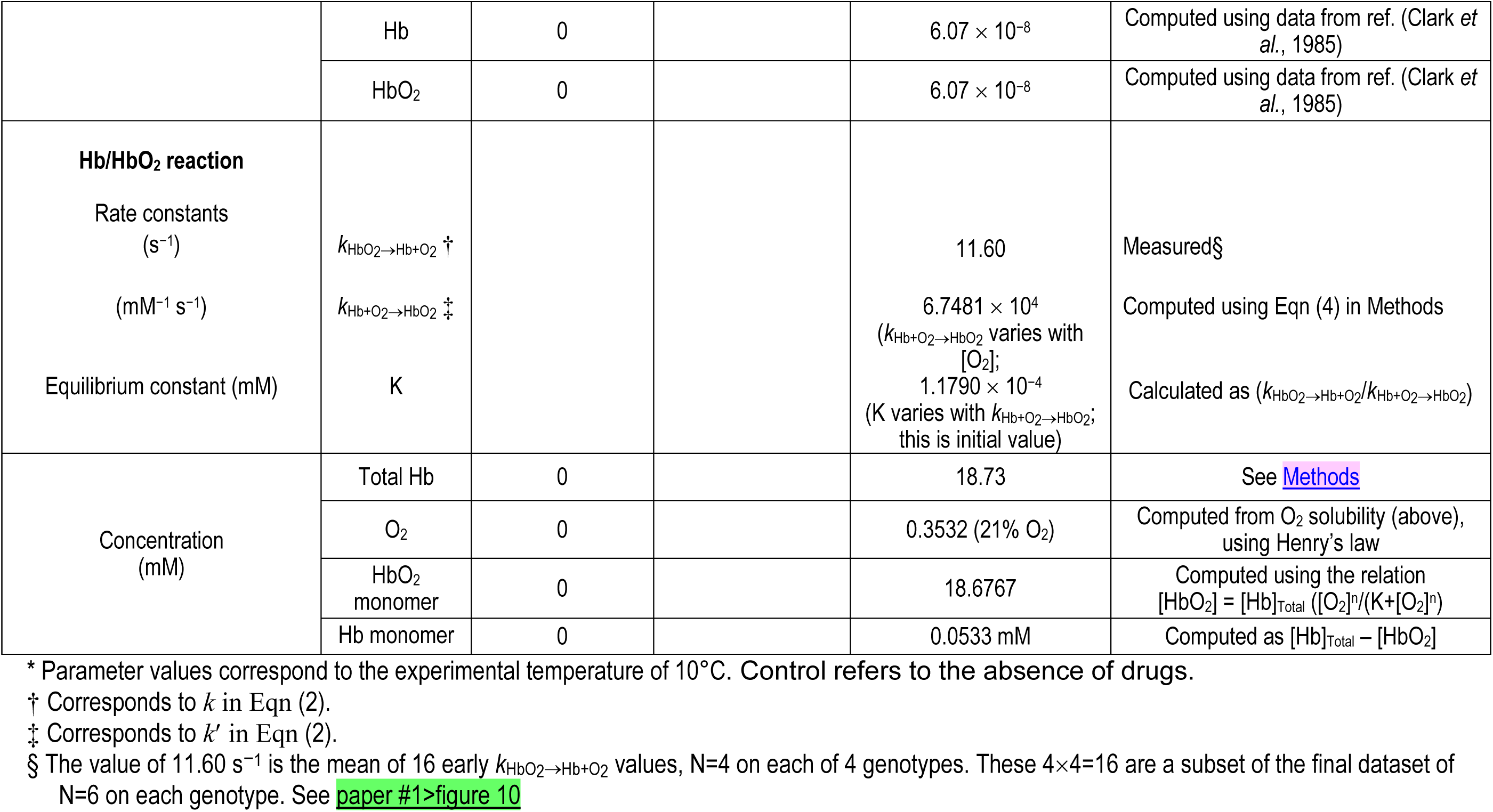
Parameter values used in mathematical simulations of control RBCs from WT mice *.

**Table 2.** Genotype-specific parameter values used in mathematical simulations of control RBCs in Figure 5.

| Parameter \ Mouse | WT | AQP1-KO | RhAG-KO | dKO | Reference |
| --- | --- | --- | --- | --- | --- |
| MCV (fl) | 47.9 | 49.5 | 51.0 | 49.8 | From paper #2>table 4 |
| $\emptyset_{\text{Major}}$, major diameter of RBC ($\mu\text{m}$) | 6.80 | 6.69 | 6.53 | 6.55 | From paper #2>table 3<br>(We assume that $OD_{\text{Torus}} = \emptyset_{\text{Major}}$) |
| $r_{\text{Torus}}$, minor radius of torus ($\mu\text{m}$) | 1.01 | 1.04 | 1.09 | 1.07 | Computed from Eqn (5) |
| $R_{\text{Torus}}$, major radius of torus ($\mu\text{m}$) | 2.39 | 2.31 | 2.18 | 2.21 | $R_{\text{Torus}} = (OD_{\text{Torus}})/2 - r_{\text{Torus}}$<br>Torus volume (computed from $r_{\text{Torus}}$ & $R_{\text{Torus}}$) = MCV |
| $\emptyset_{\text{Sphere}}$, diameter of equivalent sphere ($\mu\text{m}$) | 2.02 | 2.08 | 2.18 | 2.14 | $\emptyset_{\text{Sphere}} = 2 \times r_{\text{Torus}}$ |
| MCH, mean corpuscular hemoglobin (pg) | 14.4 | 14.1 | 14.7 | 14.5 | From paper #2>table 4 |
| MCHC, mean corpuscular hemoglobin concentration (g/dl) | 30.1 | 28.6 | 28.7 | 29.4 | From paper #2>table 4<br>MCHC = MCH/MCV |
| $[\text{Hb}_{\text{M,Total}}]_i$, total concentration of Hb monomers (mM) | 18.73 | 17.71 | 17.87 | 18.16 | See Methods > Mathematical modeling and simulations > Calculation of Hb content |
| Initial $[\text{Hb}_\text{M}\text{O}_2]_i$, concentration of $\text{O}_2$-bound monomers (mM) | 18.6767 | 17.6596 | 17.8191 | 18.1083 | Computed using the relation<br>$[\text{Hb}_\text{M}\text{O}_2] = [\text{Hb}_{\text{M,Total}}] \cdot ([\text{O}_2]^n / (K + [\text{O}_2]^n))$ |
| Initial $[\text{Hb}_\text{M}]_i$, concentration of $\text{O}_2$-free monomers (mM) | 0.0533 | 0.0504 | 0.0509 | 0.0517 | Computed as $[\text{Hb}_{\text{M,Total}}] - [\text{Hb}_\text{M}\text{O}_2]$ |
| $k_{\text{HbO}_2 \rightarrow \text{Hb} + \text{O}_2}$ | 11.60 | 12.31 | 12.03 | 10.44 | See paper #1>figure 10 |

\*Values not listed here, but necessary for the mathematical simulations, are in Table 1, which also repeats four values in the “WT” column of this table (i.e., $\varnothing_{\text{Sphere}}$ , $[\text{Hb}_{\text{M,Total}}]_{\text{i}}$ , initial $[\text{Hb}_{\text{M}}\text{O}_2]_{\text{i}}$ , Initial $[\text{Hb}_{\text{M}}]_{\text{i}}$ ). dKO, double knockout (i.e., genotype *Aqp1*<sup>-/-</sup>*Rhag*<sup>-/-</sup>). The parameters in Table 2 that establish the genotype-specific conditions for each simulation are: (1) the RBC thickness, which we compute from the measured $\varnothing_{\text{Major}}$ and MCV, and (2) the total Hb concentration, which we compute from measured MCV and MCH. Thus, the input parameters for the four simulations that mimic RBCs from WT, AQP1-KO, RhAG-KO and dKO mice are all the same (including $P_{\text{M,O}_2}$ ) with the exception of the RBC thickness and total Hb concentration, which are specific to each mouse genotype.

We solve the system of the three reaction-diffusion equations (i.e., Eqn (1) for the solutes O_2_, Hb, and HbO_2_) using the method of lines and the approach described by Somersalo et al for the numerical implementation of the diffusion component (Somersalo *et al*., 2012). We solve the resulting system of time-dependent ordinary differential equations in Matlab R2015a using the stiff solver ode15s with AbsTol=RelTol=1e–12. The Matlab code is based on the original implementation of Somersalo et al.

#### Parameter values

Table 1 lists the parameter values used in the simulation for RBCs from WT mice. Table 2 lists the parameters that we measured in the present study, specific for each of the 4 genotypes; we use these values for computing values for Figure 5. All parameter values correspond to a temperature of 10°C, the temperature at which we perform the stopped-flow experiments (see <u>paper #1</u>^16^). For parameter values not available at the temperature of 10°C, we derive them using Arrhenius equation and values at 25°C and 37°C from references indicated in Table 1. We use this approach for calculating the diffusion coefficient of HbO_2_ and HbO_2_ in the bulk ICF.

For the diffusion coefficient of O_2_ in the bulk ICF, the commonly used value (adjusted for 10°C using the Arrhenius equation, as described above) is 5.09 × 10^−6^ cm^2^ s^−1^ (Clark *et al*., 1985). More recently, Richardson et al (Richardson *et al*., 2020) obtained the much lower value of 0.45907 × 10^−6^ cm^2^ s^−1^. However, we are concerned that this latter value is an underestimate (see <u>Discussion</u>)^17^. We chose to use the arithmetic mean of the commonly used value of 5.09 × 10^−6^ cm^2^ s^−1^ and the more recent value of 0.45907 × 10^−6^ cm^2^ s^−1^, namely 2.7745 × 10^−6^ cm^2^ s^−1^ (i.e., somewhat more than half the value reported by Clark *et al*., 1985; see Table 1).

Because the molecular weight of O_2_ is much less than the molecular weight of Hb, and because the diffusion coefficient mainly depends on molecular weight, we assume that the diffusion coefficients of monomeric HbO_2_ and Hb are the same (Popel, 1989). We assume that the permeability of the RBC plasma membrane to O_2_ is 0.15 cm s^−1^, the value estimated by Endeward et al (Endeward *et al*., 2006) for the permeability of the RBC plasma membrane to CO_2_.

#### Simulation of time course of HbO2 deoxygenation, and estimation of *k*HbO2

Using the reaction-diffusion model, we simulate the time course of the decline of HbO_2_ saturation (i.e., HbSat = [HbO_2_]/([Hb]+[HbO_2_]) vs. time, employing hematological and morphological data that we gathered for RBCs from WT mice (see Table 1). Because neither the actual nor the simulated time courses of HbO_2_ desaturation are precisely exponential, we compute *k*_HbO2_ from simulations in a manner analogous to that for our physiological data (see <u>paper #1</u>^18^): we determine the time for the simulated HbSat to fall to 1/e ≅ 37% of its initial value (i.e., τ_Quasi_ ≅ *t*_37_), and then compute *k*_HbO2_ as 1/*t*_37_.

### Data availability

The data supporting the findings of this study are available within the paper. Any further relevant data are available from the corresponding author upon reasonable request.

## Results

The overarching hypothesis of papers #1 and #2 is that AQP1, RhAG, and at least one other RBC membrane protein contribute in a major way to the O_2_ permeability of murine RBC membranes.

From a technical perspective, the purpose of this third paper is to develop a reaction-diffusion model—our shorthand for this macroscopic mathematical model is MMM—that is applicable to RBCs in the stopped-flow studies of paper #1. Because our MMM incorporates Fick’s first law of diffusion to describe transmembrane O_2_ efflux (see <u>Methods</u>)^19^ we can relate the input parameter *P*_M,O2_—incorporated into Eqns (6) and (7)—to the output parameter MMM-*k*_HbO2_. As noted in the Introduction, in our simulations, we do not adjust myriad parameter values to match physiological data, but rather use the model to predict—from first principles—the time course of HbO_2_ desaturation, which yields MMM-*k*_HbO2_. The input parameters for the simulations are physical constants and values obtained from the literature (see Table 1) as well as values obtained in automated-hematology studies and imaging flow cytometry in papers #1 & #2 (see Table 2). The latter—which include MCV, MCH, and Ø_Major_—allow us to determine values for RBC thickness and total Hb concentration and thereby perform simulations that mimic as closely as possible the conditions of our physiological experiments.

From a philosophical perspective, our purpose in the present paper is to use simulations based on the MMM to make a series of estimates of *P*_M,O2_ that generate a series of MMM-*k*_HbO2_ values that allow us to hone in on our best estimate of *P*_M,O2_ in a particular set of stopped-flow experiments in paper #1.

If our hypothesis is correct, RBCs from mice genetically deficient in AQP1, RhAG, or both ought to have lower *P*_M,O2_ values when compared to RBCs from WT mice. In order to test our hypothesis, we must assess *P*_M,O2_ from experimental data. In <u>paper #1</u>^20^, we use SF absorbance spectroscopy to measure *k*_HbO2_ of RBCs from mice genetically deficient in AQP1, RhAG, or both.

We find that, when compared to WTs, *k*_HbO2_ decreases by 9% in AQP1-KOs, 17% in RhAG-KOs, and 30% in dKOs. Moreover, *k*_HbO2_ decreases by 53% in dKO RBCs treated with DIDS, and by 78% in dKO RBCs treated with PCMBS. One might immediately ask why the sum 9% + 17% is not precisely 30%? The answer, aside from experimental and rounding errors, is that *P*_M,O2_—the presumed target of our genetic and pharmacological manipulations—is only one element that contributes to MMM-*k*_HbO2_ (see Figure 2A), resulting (as we shall see later) in a curvilinear dependence of *P*_M,O2_ on log(*k*_HbO2_).

In the following sections we show that, by informing the model with reasonable parameter values and internally consistent (i.e., gathered in our laboratory) hematology and flow cytometry data, our MMM model provides reasonable estimates of the time course of HbO_2_ desaturation and corresponding MMM-*k*_HbO2_. Moreover, the model confirms that decreases in the value of *k*_HbO2_ correspond to decreases in *P*_M,O2_, thereby supporting the hypothesis that membrane proteins like AQP1 and the mouse Rh complex (Rh_Cx_ = RhAG + RhD) contribute to *P*_M,O2_.

### Simulations of O2 efflux from a control RBC of WT mouse

Our first goal was to establish an in-silico experiment that would mimic O_2_ efflux from a control (Ctrl; i.e., not pretreated with inhibitors) RBC of a WT mouse (WT/Ctrl). For this simulation, our input parameters are those reported in Table 1, which includes physical constants and values obtained from the literature as well as values for RBC thickness and total Hb concentration obtained from our hematological and morphological data (i.e., MCV, MCH, Ø_Major_) of WT mice (see Table 2), first column).

Figure 4 shows results of this simulation. At the top of the figure, panels *A*, *B*, and *C* show the time courses of the intracellular (i) concentrations of free O_2_, HbO_2_, and the sum of free O_2_ + HbO_2_—integrated over the entire volume of the cytoplasm—as O_2_ diffuses out of the cell in a simulated O_2_-offloading experiment, as in <u>paper #1</u>^21^.

**Figure 4.**
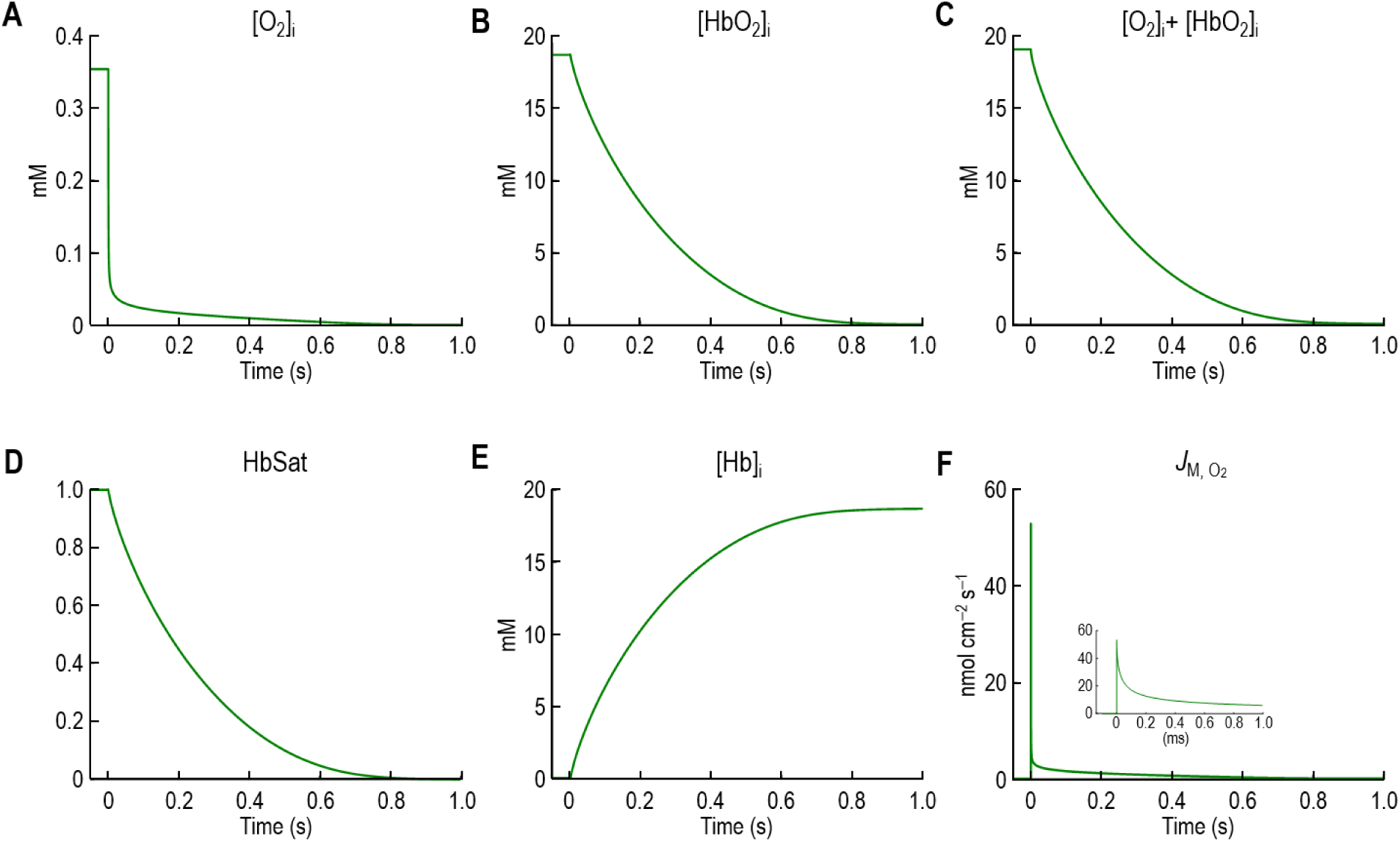
Simulated time courses of O_2_/Hb-related parameters and transmembrane O_2_ flux for cells mimicking WT/Ctrl (control/no inhibitors) For this simulation, the input parameters are those reported in Table 1. See text for details on the model and simulations. All concentrations are in mM. *A,* Concentration of free O_2_, integrated over the entire volume of cytoplasm. *B,* Concentration of oxyhemoglobin (HbO_2_) monomers, integrated over the entire volume of cytoplasm. *C,* Concentration of the sum of free O_2_ and HbO_2_ monomers, integrated over the entire volume of cytoplasm. *D,* Hemoglobin saturation (HbSat), integrated over the entire volume of cytoplasm. *E,* Concentration of free hemoglobin (Hb) monomers, integrated over the entire volume of cytoplasm. *F,* Transmembrane O_2_ flux (*J*_M,O2_). Note that the units are nmol cm^−2^ s^−1^ (rather than μmol cm^−2^ s^−1^, as in the text of the Discussion). The inset shows a magnification of the first 1 ms of the simulated *J*_M,O2_

**Figure 5.**
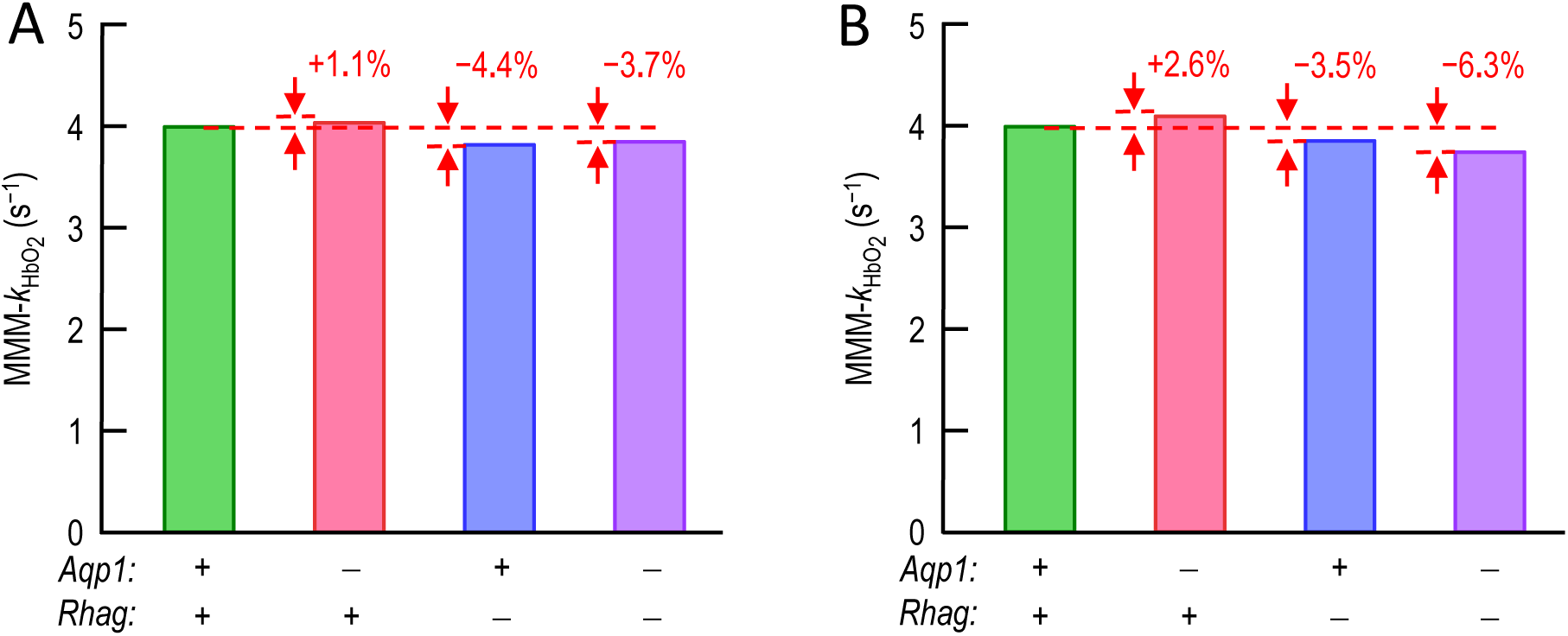
MMM-*k*_HbO2_ values predicted from simulations of RBCs from WT and KO mice Informed by hematological, morphological, and kinetic data, which provide genotype-specific information on MCV, MCH, Ø_Major_, and *k*_HbO2→Hb+O2_ (see Table 2) we perform four simulations, one per genotype, to test whether the collection of parameter changes can explain the physiologically observed decreases in shape-corrected (SC) *k*_HbO2_ values reported in figure 5*b* <u>of paper #1</u>: about –9% (AQP1-KO), –17% (RhAG-KO), and –30% (double KO). The values in red above each knockout bar indicate the % change predicted by MMM-*k*_HbO2_ vs. the observed change. Thus, the hematological/morphological changes cannot explain our observed *k*_HbO2_ data. *A, k*_HbO2_→_Hb+O2_ assumed to be 11.60 s^−1^ for all genotypes. *B, k*_HbO2_→_Hb+O2_ assumed to be specific for each genotype. Except as noted otherwise, in these simulations we used the values summarized in Table 1, including *P*_M,O2_ = 0.15 cm s^−1^. We assume that all cells are intact (analogous to a hemolysis correction), and that all cells are biconcave disks (analogous to a shape correction). Thus, the two green bars have MMM-*k*_HbO2_ values of ∼3.99 s^−1^. MMM-*k*_HbO2_, macroscopic mathematically modeled rate constant for deoxygenation of intracellular HbO_2_; *Aqp1*, the gene encoding murine aquaporin 1; *Rhag*, the gene encoding murine RhAG: +, gene present; –, gene deleted. MCV, mean corpuscular volume; MCH, mean corpuscular hemoglobin; Ø_Major_, major diameter of biconcave disk; *k*_HbO2→Hb+O2_, rate constant for deoxygenation of HbO_2_ in free solution; *P*_M,O2_, membrane permeability to O_2_.

### [O2]i

As expected, because O_2_ is leaving the cell, [O_2_]_i_ decreases (Figure 4A).

### [HbO2]i and [O2]i + [HbO2]i

At the same time as [O_2_]_i_ decreases, [HbO_2_]_i_ decreases (Figure 4B) in parallel with intracellular [O_2_]_i_ + [HbO_2_]_i_ (Figure 4C). These three time courses in Figure 4A-C reflect (1) the efflux of free O_2_; (2) the diffusion of free O_2_ toward the membrane; (3) the net reaction HbO_2_ → Hb + O_2_, which consumes HbO_2_ to produce free Hb and free O_2_ and continues until HbO_2_ is fully depleted; (4) the diffusion of HbO_2_ toward the membrane, and (5) the diffusion of free Hb away from the membrane. The time courses of [HbO_2_]_i_ and of [O_2_]_i_ + [HbO_2_]_i_ are nearly identical because [O_2_]_i_ << [HbO_2_]_i_.

At the bottom of the figure, panels *D*, *E*, and *F* show the time courses of hemoglobin saturation, [Hb]_i_, and the O_2_ efflux across the plasma membrane (*J*_M,O2_).

### HbSat

As expected, HbSat falls (Figure 4D) with a time course that is similar to that of [HbO_2_]_i_ (see Figure 4B). It is from this time course of HbSat that we calculate the MMM-*k*_HbO2_ of WT/Ctrl RBCs (see <u>Methods</u>^22^). The model predicts a MMM-*k*_HbO2_ of ∼3.99 s^−1^, which will become our “Provisional MMM-*k*_HbO2_” in our accommodation for nBCDs (see <u>below</u>^23^).

### [Hb]i

As anticipated, [Hb]_i_ increases (Figure 4*E*) with a time course that is the inverse of that of [HbO_2_]_i_.

### *J*M,O_2_

Finally, *J*_M,O2_ initially zero, spikes rapidly to a maximal value, decays rapidly to a value somewhat above zero, and finally drifts toward zero (Figure 4*F*). The *J*_M,O2_ upward spike reflects the large initial transmembrane O_2_ gradient. The rapid decay reflects O_2_ depletion near the inner surface of the membrane. The O_2_ efflux during the slower decay is supported by O_2_ that diffuses to the inner surface of the membrane from the depths of the RBC and, to a much lesser extent, the parallel diffusion and dissociation of HbO_2_.

### Predicted *k*HbO_2_ values from simulations of control RBCs from WT and KO mice, as informed by genotype-specific hematological, morphological, and kinetic data

Besides *P*_M,O2_, the MMM-*k*_HbO2_ that describes O_2_-offloading from RBCs depends on several genotype-specific parameters: Ø_Sphere_ (which in turn depends on MCV, Ø_Major_, and RBC shape), [Hb_Total_]_i_ (computed from MCHC), and *k*_HbO2→Hb+O2_ (i.e., the same as *k* in Eqn (2); determined on SF experiments on RBC hemolysates). To investigate whether the decreases in *k*_HbO2_ observed in RBCs from KOs (see <u>paper #1</u>^24^) could be explained by changes in parameters other than *P*_M,O2_, we as a first step gathered the requisite genotype-specific data in paper #1 and paper #2. We provide these parameter values in Table 2. Note the trend (from left to right) for MCV to increase in the KOs, but for MCHC to decrease. In addition, using a battery of methodologies to assess RBCs shape (blood smears, still/video images of living RBCs, and imaging flow cytometry), we find that ∼98% of control RBCs, both from WTs and dKOs, are biconcave disks (see <u>paper #2</u>^25^).

To assess the effects of the genotype-specific differences summarized in Table 2, we perform simulations in which we mimic control RBCs (i.e., not pretreated with inhibitors) from WT, AQP1-KO, RhAG-KO and dKO mice. Note that the simulation for a control WT RBC is the same one that yielded the MMM-*k*_HbO2_ of 3.99 s^−1^ in the previous section. We simulate O_2_ efflux from RBCs of AQP1-KO/Ctrl, RhAG-KO/Ctrl, and dKO/Ctrl mice using the same parameter values (including *P*_M,O2_) that we used for WT/Ctrl RBC (see Table 1) with the exception of the genotype-specific values in Table 2 for Ø_Sphere_ (mimicking RBC thickness) and [Hb_M,Total_]_i_ (a surrogate for MCHC).

Figure 5 shows a pair of MMM-*k*_HbO2_ values for each of the four genotypes. In these simulations we assume that all RBCs are intact, as they are for the hemolysis-corrected (HC) *k*_HbO2_ values in paper #1, and that all cells are BCDs, as they are for the shape-corrected (SC) *k*_HbO2_ values for SF experiments in <u>paper #1</u>^26^. In Figure 5*A*, we compute MMM-*k*_HbO2_ values using our standard *k*_HbO2→Hb+O2_ value of 11.6 s^−1^ (see Table 1), which is an average for all four genotypes. We use this standard value for all simulations here in paper #3 (including Figure 5*A*) except for those in Figure 5*B*, where we use four separate, genotype-specific *k*_HbO2→Hb+O2_ values (see Table 2).

The solid green bars in both Figure 5*A* and *B* correspond to an MMM-*k*_HbO2_ of ∼3.99 s^−1^ for a WT/Ctrl RBC (i.e., no drugs). Both panels have the same MMM-*k*_HbO2_ value because, by serendipity, the standard *k*_HbO2→Hb+O2_ is virtually the same as the genotype-specific value for WT/Ctrl RBCs.

The pink bar in Figure 5*A* simulates an AQP1-KO/Ctrl RBC; MMM-*k*_HbO2_ ≅ 4.04 s^−1^, which is a predicted 1.1% increase vs. 3.99 s^−1^, rather than the 9% decrease observed in paper #1. In Figure 5*B*, MMM-*k*_HbO2_ is ∼4.10 s^−1^, which represents an even larger increase, 2.6%.

The blue bar in Figure 5*A* simulates a RhAG-KO/Ctrl RBC; its MMM-*k*_HbO2_ of 3.82 s^−1^ represents a predicted decrease of 4.4%, rather than the 17% decrease observed in paper #1. The corresponding MMM-*k*_HbO2_ in Figure 5*B*, is ∼3.85 s^−1^, which predicts a somewhat smaller decrease of 3.5%.

Finally, the purple bar in Figure 5*A* simulates a dKO/Ctrl RBC; its MMM-*k*_HbO2_ of 3.85 s^−1^ is a decrease of only 3.7% vs. WT/Ctrl, rather than the ∼30% decrease in paper #1. In Figure 5*B*, MMM-*k*_HbO2_ is ∼3.74 s^−1^, which predicts a decrease of 6.3%, which is still far smaller than what we observed in paper #1.

Thus, we conclude that the genotype-specific changes reported in paper #1 and paper #2— and summarized here in Table 2—cannot explain the observed KO-induced decreases in SC-*k*_HbO2_ in paper #1. We intuit that the effects of anti-parallel increases in MCV and decreases in MCHC tend to cancel, and thereby tend to minimize the impact on MMM-*k*_HbO2_ values. We examine this possibility in the sensitivity analyses, <u>below</u>^27^. We conclude that KO-induced variations in RBC dimensions, [Hb_Total_]_i_, and *k*_HbO2→ Hb+O2_ cannot account for the KO-induced decreases in SC-*k*_HbO2_ observed in paper #1.

### Predicted dependence of O2 offloading (*k*HbO2) on the O2 permeability of RBC membrane (*P*M,O_2_)

In the previous sections, we show that our MMM—with a fixed *P*_M,O2_ and genotype-specific values for other parameters—can mimic the time course of O_2_ offloading (i.e., MMM-*k*_HbO2_ ≅ SC-*k*_HbO2_) for a control RBC from a WT mouse, but not for control RBCs from any of our three KO groups. Clearly, we must invoke some other parameter to account for the control and KO data in paper #1 and, by extension, the paper-#1 data on inhibitors and KO+inhibitors. We hypothesize that this other parameter is *P*_M,O2_. (In the case of drug-treated RBCs, we will apply a shape correction, described below, to accommodate the appearance of nBCDs, i.e., poikilocytes).

Having established an in-silico approach for WT/Ctrl experiments, we can now employ the model to answer the two key questions that are at the core of our investigation:

1. What is the dependence of MMM-*k*_HbO2_ on *P*_M,O2_? This will help us understand the extent to which observed decreases in SC-*k*_HbO2_ in paper #1 correspond to decreases in *P*_M,O2_?
2. Can we use physiological *k*_HbO2_ measurements to make predictions about *P*_M,O2_?

To answer these questions, we employ the model that mimics a control WT RBC (i.e., input parameter values listed in Table 1) and perform a series of more than 70 simulations in which we systematically vary *P*_M,O2_ while holding all other parameters constant. Throughout, we assume that the RBCs are biconcave disks so that we need not accommodate for shape changes. Thus, by definition, the MMM-*k*_HbO2_ is analogous to the SC-*k*_HbO2_. The continuous curve in Figure 6 is a semi-logarithmic plot showing the dependence of MMM-*k*_HbO2_ (y-axis) on *P*_M,O2_ (x-axis, scaled logarithmically). The shape of this curve depends on the parameters in Table 1, and thus could vary with each strain of mouse, and each species (see <u>Discussion</u>^28^).

**Figure 6.**
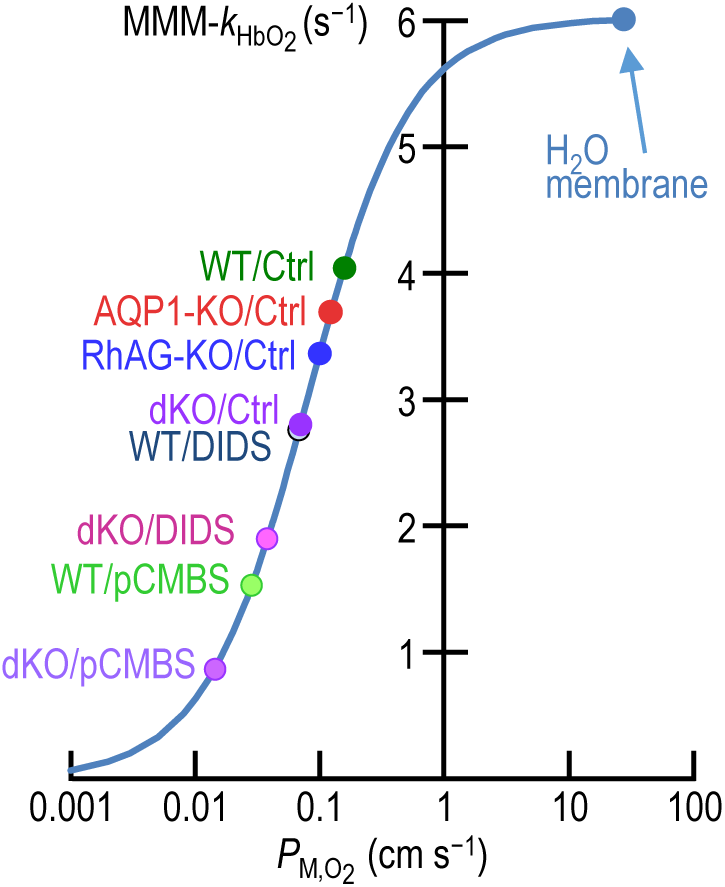
Predicted dependence of MMM-*k*_HbO2_ on *P*_M,O2_ Using our macroscopic mathematical model (MMM), we systematically varied membrane O_2_ permeability (*P*_M,O2_; logarithmic x-axis) to compute the corresponding rate constant for deoxygenation of HbO_2_ (*k*_HbO2_; linear y-axis). Except for *P*_M,O2_, we used the values summarized in Table 1 for these 70^+^ simulations. The resulting sigmoidal curve represents all possible combinations of *P*_M,O2_ and *k*_HbO2_ for *P*_M,O2_ > 0.001 cm s^−1^. The point labeled “H_2_O membrane” is the result of a simulation in which we assumed that the diffusion constant of O_2_ in the plasma membrane is the same as in water; it has the coordinates (*P*_M,O2_ = 22.63 cm s−^1^, *k*_HbO2_ = 5.9953 s−^1^). The point labeled WT/Ctrl (i.e., RBCs from wild-type mice in the absence of drugs) has the coordinates (*P*_M,O2_ = 0.1546 cm s^−1^, *k*_HbO2_ = 4.0358 s^−1^), and represents the shape-corrected value obtained by the linear-combination procedure (steps #11 through #22 in Figure 1). The other labeled points also represent shape-corrected values, as summarized in Table 3. Note that the dKO/Ctrl and the WT/DIDS points nearly overlie each other.

**Table 3.** Parameter values predicted by macroscopic mathematical model.

| Protocol<br>Parameter | WT/<br>Ctrl | AQP1-KO/Ctrl | RhAG-KO/Ctrl | dKO/<br>Ctrl | WT/<br>pCMBS | WT/<br>DIDS | dKO/<br>pCMBS | dKO/<br>DIDS |
| --- | --- | --- | --- | --- | --- | --- | --- | --- |
| SC-$P_{M,O_2}$ (cm s<sup>-1</sup>) | 0.1546 | 0.1213 | 0.0983 | 0.0693 | 0.0281 | 0.0673 | 0.0143 | 0.0376 |
| SC-$k_{HbO_2}$ (s<sup>-1</sup>) | 4.0358 | 3.6828 | 3.3582 | 2.7988 | 1.5282 | 2.7533 | 0.8687 | 1.8937 |
| ↓ SC-$P_{M,O_2}$ vs. WT/Ctrl | 0% | 21.56% | 36.43% | 55.20% | 81.80% | 56.47% | 90.72% | 75.71% |
| ↓ SC-$k_{HbO_2}$ vs. WT/Ctrl | 0% | 8.85% | 16.53% | 30.47% | 61.24% | 31.25% | 77.70% | 52.98% |

The SC- $P_{M,O_2}$ (via the MMM) yields the SC- $k_{HbO_2}$ .
Ctrl, control conditions (i.e., no DIDS or pCMBS); MMM, macroscopic mathematical model; SC, shape corrected.

The point at the extreme upper-right—labeled “H_2_O membrane”—represents the hypothetical situation in which we assume that the plasma membrane—as is implicit in Krogh’s calculations—is a thin film of H_2_O of equivalent thickness. That is, for the plasma membrane, we assume that the diffusion constant *D*_O2_ has the same value as in free water, namely 1.3313 × 10^−5^ cm^2^ s^−1^, which translates to a *P*_M,O2_ of ∼22.63 cm s^−1^. In our simulations, this value generates a MMM-*k*_HbO2_ of ∼6.00 s^−1^. For the given geometric, hematologic, and kinetic parameters, this point represents the maximum-possible values for *P*_M,O2_ and thus the maximum-possible rate of O_2_ offloading.

What is the position of a WT/Ctrl RBC on this curve? Note that (1) WT/Ctrl RBCs have an nBCD prevalence of 1.41% (see <u>paper #2</u>^29^) and (2) nBCDs (because of their larger spherical diameter; see paper #2^29^) are expected to have a lower O_2_-offloading rate that BCDs. The MMM-*k*_HbO2_ of 3.99 s^−1^—which we have already generated using *P*_M,O2_ = 0.15 cm s^−1^—in fact pertains not to a pure population of BCDs, but rather to a mixture of 98.59% BCDs and 1.41% nBCDs. Our accommodation for nBCDs presented <u>below</u>^30^) leads to the conclusions that (1) the actual *k*_HbO2_ for the BCD component is ∼4.04 s^−1^ vs. 3.99 s^−1^ for the mixture, and (2) the actual *P*_M,O2_ for both BCDs and nBCDs is 0.1546 cm s^−1^, which is ∼3.1% higher than our provisional value of 0.15 cm s^−1^. Thus, the point labeled “WT/Ctrl” in Figure 6 has the shape-corrected coordinates (*P*_M,O2_ = 0.1546 cm s^−1^, *k*_HbO2_ = 4.0358 s^−1^) and is our best estimate of a pure population of BCDs.

Note that the *P*_M,O2_ of 0.1546 cm s^−1^ for WT/Ctrl BCDs is only ∼0.7% as large (or ∼1/150) as the hypothetical *P*_M,O2_ of Krogh’s pure-water membrane. In other words, the RBC membrane offers 150-fold more resistance to O_2_ diffusion than would a film of H_2_O.

Besides WT/Ctrl, each of the seven other labeled points in Figure 6 corresponds to one of the other experimental conditions. For each point, we determine the y-axis value (MMM-*k*_HbO2_) by performing the shape correction described below^30^ (steps #13 – #22 in Figure 1), based on experimentally determined parameter values. We determine the x-axis value (*P*_M,O2_) by interpolation (step #19 in Figure 1), thereby placing the point on the sigmoidal curve. Note that the dKO/Ctrl and the WT/DIDS points nearly overlap.

Table 3 summarizes the key numerical values corresponding to the eight experimental points in Figure 6. Our standard condition is WT/Ctrl (upper-left corner of table), for which the shape-corrected values are SC-*P*_M,O2_ = 4.0358 s^−1^ and SC-*P*_M,O2_ = 0.1546 cm s^−1^. Compared to these standard SC values, the AQP1-KO/Ctrl has an SC-*P*_M,O2_ that is 21.56% lower, and an SC-*k*_HbO2_ that is 8.85% lower—and so on for the other conditions. Note that the sum of the computed % decreases in *P*_M,O2_ for AQP1-KO/Ctrl (∼22%) and RhAG-KO/Ctrl (∼36%) is nearly equal to the computed *P*_M,O2_ % decrease for the dKO/Ctrl (∼55%). The largest % decrease in *P*_M,O2_ occurs with dKO/pCMBS (∼91%). The effects of the membrane-protein knockouts and inhibitors on *P*_M,O2_—along with the proteomics data in <u>paper #2</u>^31^—supports the hypothesis that the vast majority of O_2_ moves through channels in RBC plasma membrane: AQP1, Rh_Cx_, and at least one additional channel that is blocked by pCMBS. In other words, under our experimental conditions, “Overton’s rule” applies, at most, to ∼9% of the O_2_ traffic across the murine RBC membrane.

### Predicted effects of dKO/pCMBS on the kinetics of parameters related to intracellular O2 and Hb

In Figure 4*A-F*, we examined the simulated time courses of six parameters for control (drug-free) WT RBCs: [O_2_]_i_, [HbO_2_]_i_, [O_2_]_i_ + [HbO_2_]_i_, HbSat, [Hb]_i_, and *J*_M,O2_. In Figure 7*A-F*, we extend this WT/Ctrl analysis (green curves) by simulating the slowest O_2_ offloading protocol observed in <u>paper #1</u>^32^: a dKO RBC pre-treated with pCMBS (purple curves).

**Figure 7.**
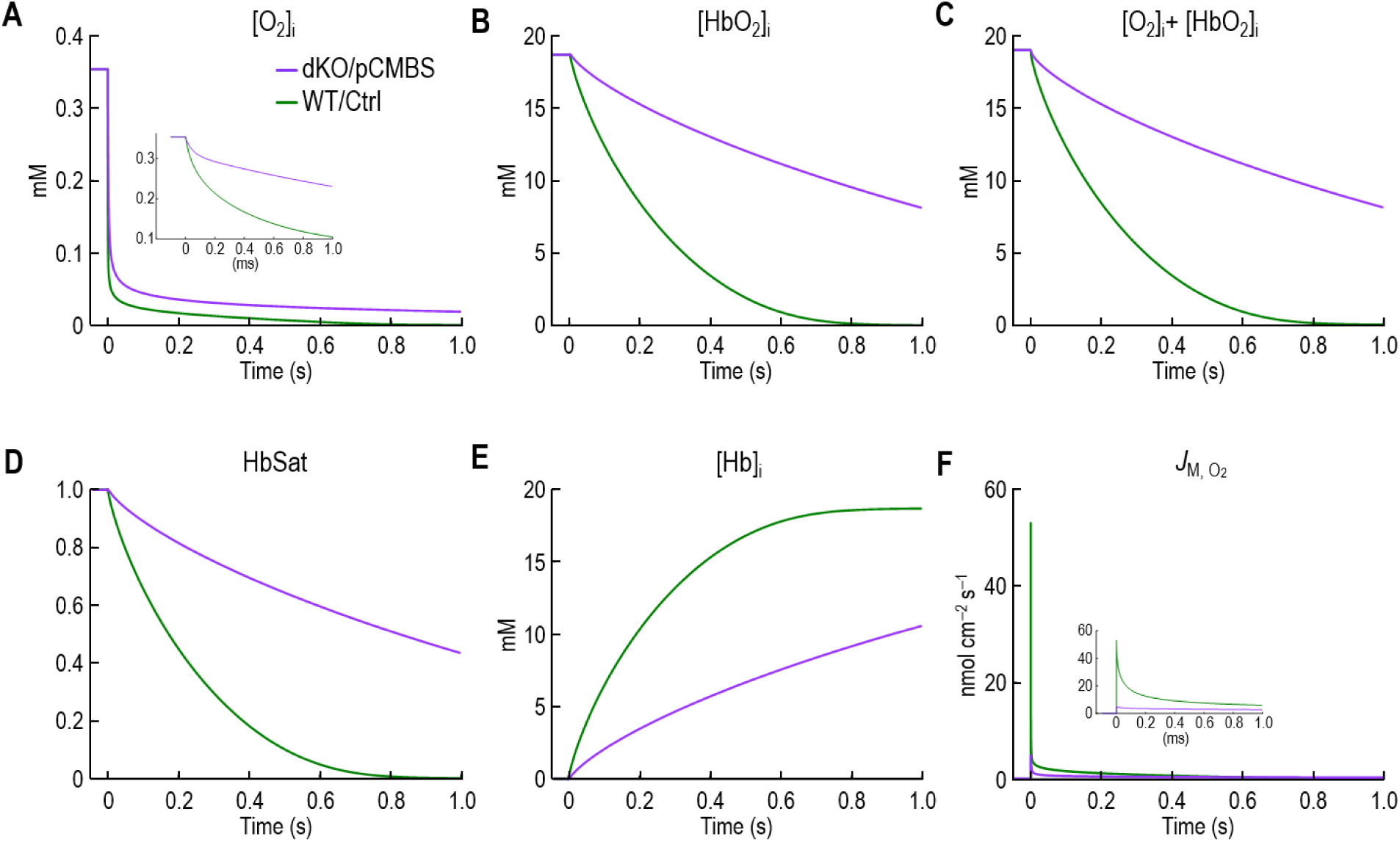
Simulated time courses of O_2_/Hb-related parameters and transmembrane O_2_ flux for cells mimicking WT/Ctrl (control/no inhibitors) vs. dKO/pCMBS We extend the analysis of the WT/Ctrl condition (untreated RBCs from wild-type mice; green curves) shown in Figure 4 by adding the simulations of the slowest O_2_-offloading protocol observed in <u>paper #1 (i.e.,</u> figure 5*b*<u>, light purple bar)</u>: the dKO/pCMBS condition (i.e., RBCs pre-treated with pCMBS; purple curves). For the WT/Ctrl simulation, the input parameters are those reported in Table 1. For the dKO/pCMBS simulation, the input parameters are the same, with the exception of *P*_M,O2_ = 0.01365 cm s^−1^. See text for details on the model and simulations. All concentrations are in mM. *A,* Concentration of free O_2_, integrated over the entire volume of cytoplasm. The inset shows a magnification of the first 1 ms of the simulated [O_2_]_i_. *B,* Concentration of oxyhemoglobin (HbO_2_) monomers, integrated over the entire volume of cytoplasm. *C,* Concentration of the sum of free O_2_ and HbO_2_ monomer, integrated over the entire volume of cytoplasm. *D,* Hemoglobin saturation (HbSat) integrated over the entire volume of cytoplasm. *E,* Concentration of free hemoglobin (Hb), integrated over the entire volume of cytoplasm. *F,* Transmembrane O_2_ flux (*J*_M,O2_). Note that the units are nmol cm^−2^ s^−1^ (rather than μmol cm^−2^ s^−1^, as in the text of the Discussion). The inset shows a magnification of the first 1 ms of the simulated *J*_M,O2_

On the time scale of the main part of Figure 7*A*, the initial rates of decline for [O_2_]_i_ appear to be nearly the same for dKO/pCMBS and WT/Ctrl. The inset—a blow-up of the initial 1 ms of the simulation—shows that, in fact, the time course for dKO/pCMBS is much slower than for WT/Ctrl, reflecting a *P*_M,O2_ that is reduced by ∼91%.

For [HbO_2_] in Figure 7*B*, the time course of the decline for dKO/pCMBS is markedly slower than for WT/Ctrl.

The time courses of [O_2_]_i_ + [HbO_2_]_i_ in Figure 7*C* are nearly the same as for [HbO_2_] alone in Figure 7*B* because, of the total O_2_ in the RBC, ∼98% is bound to Hb as HbO_2_.

The time courses of HbSat in Figure 7*D*, when appropriately scaled, are nearly the same as for [HbO_2_]_i_ in Figure 7*B* because [HbO_2_]_i_ is the numerator in the calculation of HbSat.

Because the sum [Hb]_i_ + [HbO_2_]_i_ is constant, the time courses of [Hb]_i_ in Figure 7*E* is the inverse of [HbO_2_]_i_ in Figure 7*B*.

Finally, in Figure 7*F*, the transmembrane flux *J*_M,O2_ is initially ∼10-fold higher for WT/Ctrl than for dKO/pCMBS. For dKO/pCMBS, the *J*_M,O2_ time course has a similar shape, but is scaled down by 91%, reflecting the reduced *P*_M,O2_.

### Mathematical simulations exploring the predicted sensitivity of *k*HbO2 to eight key kinetic and geometric parameters

To understand better the dependence of O_2_ offloading on a range of parameters in Table 1, we performed a sensitivity analysis, studying eight key parameters. See Figure 2 for an overview of the model. Our approach was to start with the parameter values in Table 1; that is, with the provisional value *P*_M,O2_ = 0.15 cm s^−1^, our simulation yields the provisional MMM-*k*_HbO2_ = 3.99 s^−1^. Next, as described in the following five sub-sections, we computed *k*_HbO2_ as we systematically varied, one at a time, each of the following: *k*_HbO2→Hb+O2_ in Figure 8, [Hb_Total_]_i_ in Figure 9, each of the four diffusion constants in Figure 10*A–D*, ℓ_EUF_ in Figure 11, and Ø_Sphere_ in Figure 12.

**Figure 8.**
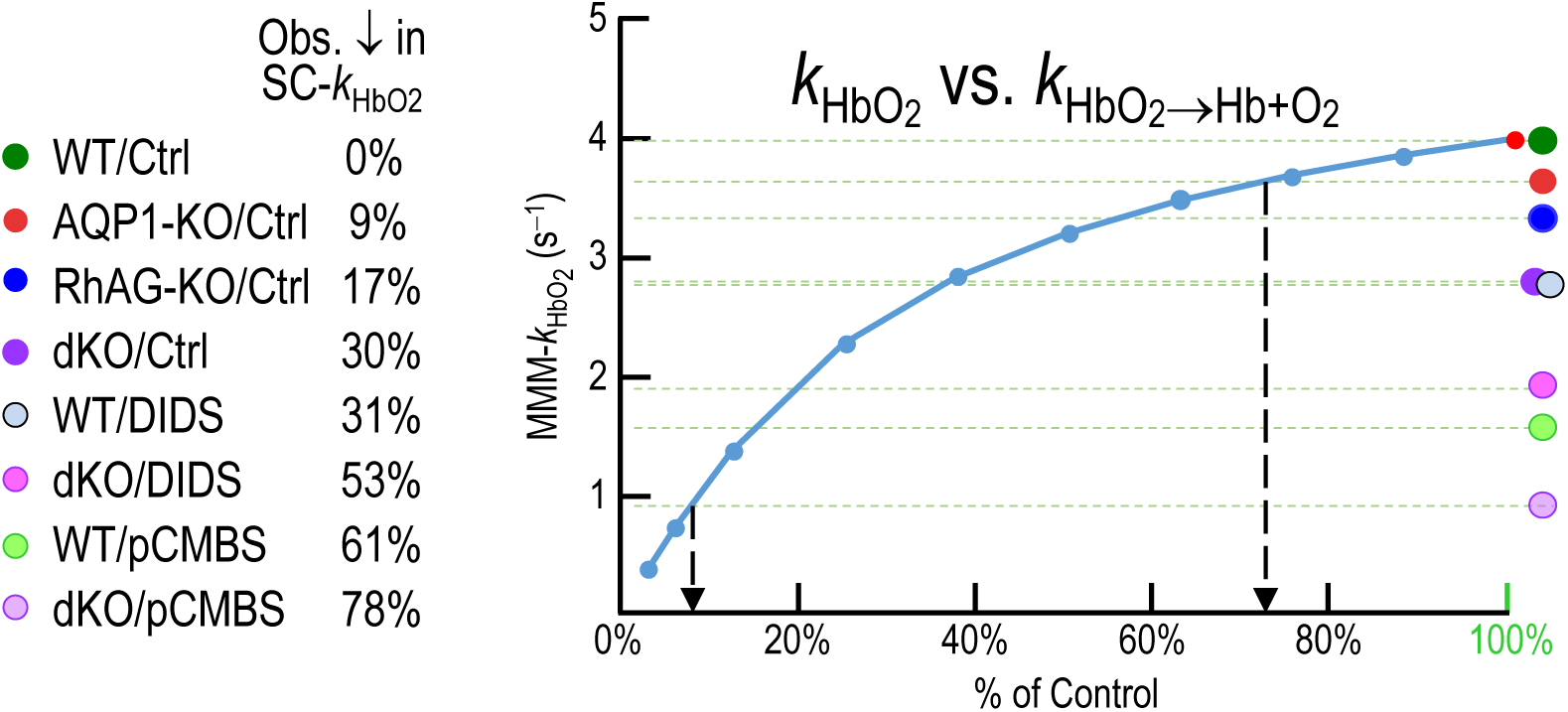
Mathematical simulations exploring the predicted sensitivity of *k*_HbO2_ to changes in the rate constant of the reaction HbO_2_ → Hb + O_2_. Using our macroscopic mathematical model (MMM), we systematically varied the rate constant for deoxygenation of HbO_2_ in free solutions: HbO_2_ → Hb + O_2_ (*k*_HbO2→Hb+O2_) to compute the corresponding the rate constant for deoxygenation of HbO_2_ in intact RBCs (*k*_HbO2_). We computed the MMM-*k*_HbO2_ corresponding to the red dot—which corresponds to 100% of our standard *k*_HbO2_→_Hb+O2_—by using the parameter values that come from Table 1. This is our standard condition for RBCs from wild-type mice in the absence of drugs (WT/Ctrl), and also is represented by the dark green circle and associated horizontal dashed green line. We performed nine additional simulations (blue dots) by decreasing *k*_HbO2_→_Hb+O2_ by the indicated percentages, keeping all other parameter values constant. The seven other colored circles and associated horizontal dashed lines indicate the MMM-*k*_HbO2_ values, relative to WT/Ctrl, for each of the other 7 experimental conditions. As summarized in the legend on the left, for each condition, we multiplied the standard MMM-*k*_HbO2_ (i.e., 3.99 s^−1^) by the shape-corrected decrease in the experimentally observed *k*_HbO2_ (e.g., ∼9% for AQP1-KO/Ctrl cells) to obtain the y-axis coordinate for that condition (e.g., 3.64 s^−1^ for AQP1-KO/Ctrl). These % decreases are also listed in the last row of Table 3. The left downward arrow (tail on dKO/pCMBS) indicates the % decrease necessary to account for the effect of this condition on MMM-*k*_HbO2_. The right downward arrow is the same for AQP1-KO/Ctrl.

**Figure 9.**
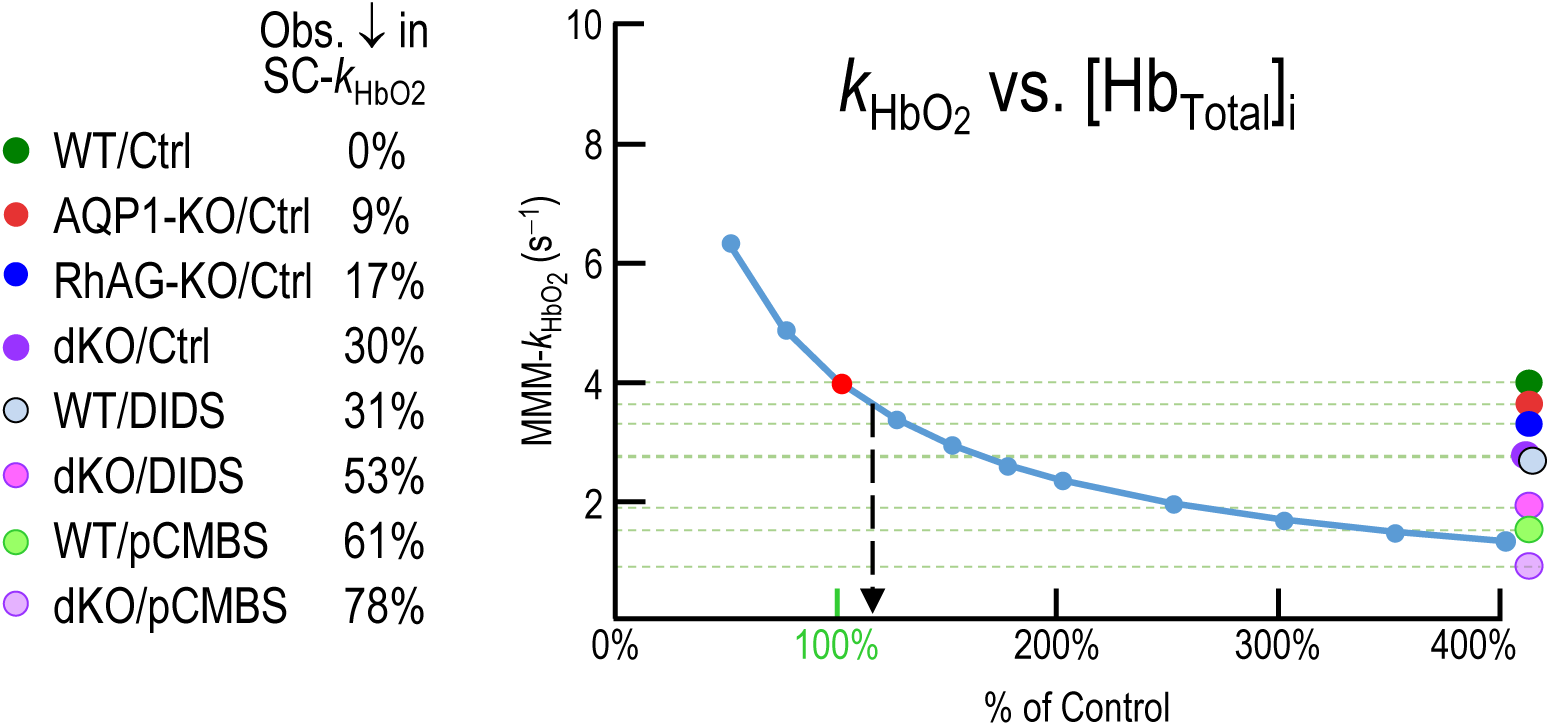
Mathematical simulations exploring the predicted sensitivity of *k*_HbO2_ to changes in the total intracellular Hb concentration Using our macroscopic mathematical model (MMM), we systematically varied the total intracellular Hb concentration, [Hb_Total_]_i_, to compute the corresponding rate constant for deoxygenation of HbO_2_ in intact RBCs (*k*_HbO2_). We computed the MMM-*k*_HbO2_ corresponding to the red dot—which corresponds to 100% of our standard [Hb_Total_]_i_—by using the parameter values that come from Table 1. This is our standard condition for RBCs from wild-type mice in the absence of drugs (WT/Ctrl), and also is represented by the dark green circle and associated horizontal dashed green line. We performed ten additional simulations (blue dots) by decreasing [Hb_Total_]_i_ by the indicated percentages, keeping all other parameter values constant. The seven other colored circles and associated horizontal dashed lines indicate the MMM-*k*_HbO2_ values, relative to WT/Ctrl, for each of the other 7 experimental conditions. As summarized in the legend on the left, for each condition, we multiplied the standard MMM-*k*_HbO2_ (i.e., 3.99 s^−1^) by the shape-corrected decrease in the experimentally observed *k*_HbO2_ (e.g., ∼9% for AQP1-KO/Ctrl cells) to obtain the y-axis coordinate for that condition (e.g., 3.64 s^−1^ for AQP1-KO/Ctrl). These % decreases are also listed in the last row of Table 3. The downward arrow (tail on AQP1-KO/Ctrl) indicates the % decrease necessary to account for the effect of this condition on MMM-*k*_HbO2_.

**Figure 10.**
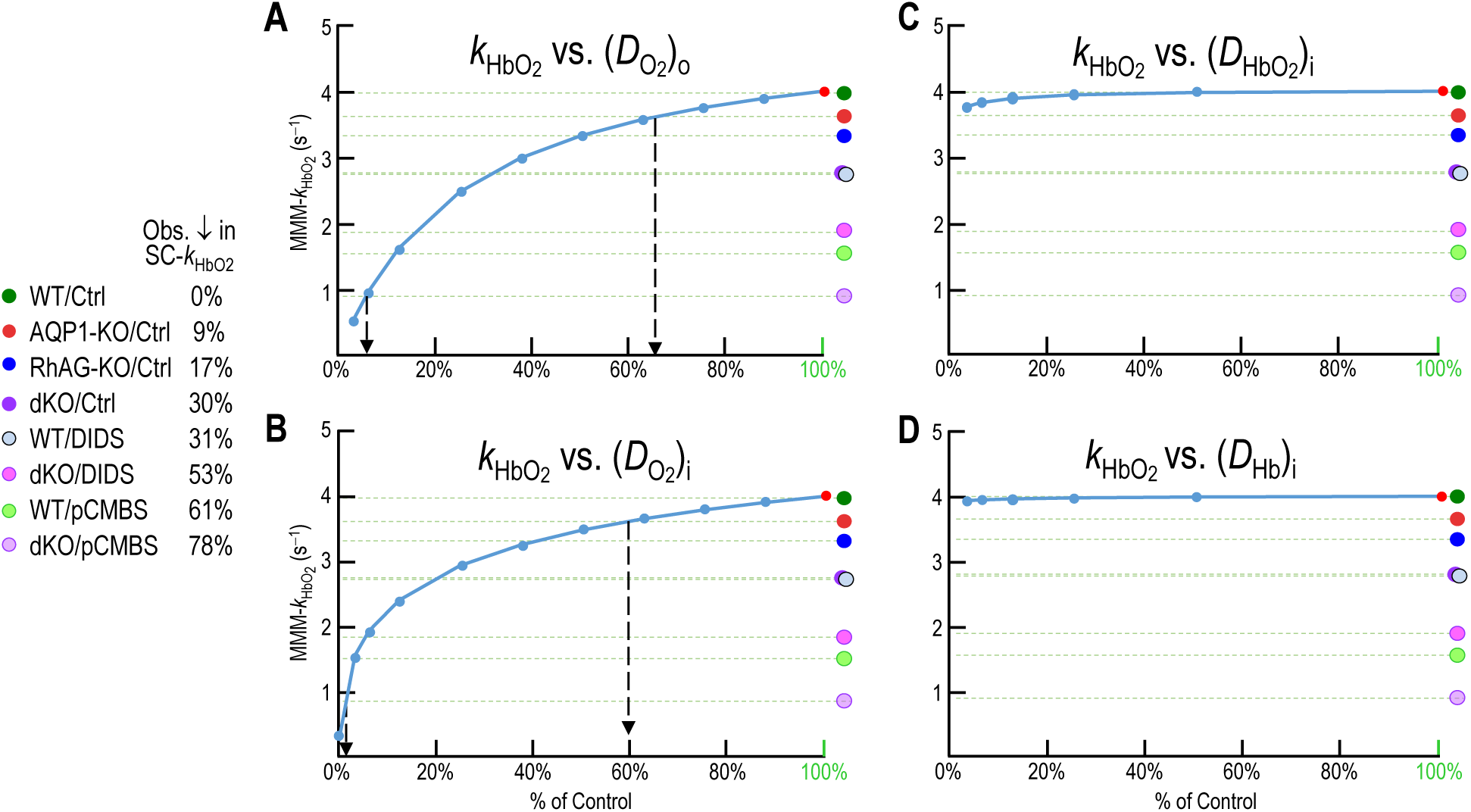
Mathematical simulations exploring the predicted sensitivity of *k*_HbO2_ to changes in diffusion constants of solutes Using our macroscopic mathematical model (MMM), we systematically varied the four relevant diffusion constants (*D*), one at a time, to compute the corresponding rate constant for deoxygenation of HbO_2_ in intact RBCs (*k*_HbO2_). These four diffusion constants are: *D* for extracellular O_2_—(*D*_O2_)_o_ in panel *A*; and the three intracellular diffusion constants for the components of the equilibrium HbO_2_ ^⇒^ Hb + O_2_, namely, (*D*_O2_)_i_ in panel *B*, (*D*_HbO2_)_i_ in panel C, and (*D*_Hb_)_i_ in panel *D*. We computed the MMM-*k*_HbO2_ corresponding to the red dot—which corresponds to 100% of the value of the varied parameter—by using the parameter values that come from Table 1. This is our standard condition for RBCs from wild-type mice in the absence of drugs (WT/Ctrl), and also is represented by the dark green circle and associated horizontal dashed green line. We performed additional simulations (blue dots) by decreasing the varied parameter by the indicated percentages, keeping all other parameter values constant. The seven other colored circles and associated horizontal dashed lines indicate the MMM-*k*_HbO2_ values, relative to WT/Ctrl, for each of the other 7 experimental conditions. As summarized in the legend on the left, for each condition, we multiplied the standard MMM-*k*_HbO2_ (i.e., 3.99 s^−1^) by the shape-corrected decrease in the experimentally observed *k*_HbO2_ (e.g., ∼9% for AQP1-KO/Ctrl cells) to obtain the y-axis coordinate for that condition (e.g., 3.64 s^−1^ for AQP1-KO/Ctrl). These % decreases are also listed in the last row of Table 3. The downward arrows (tails on AQP1-KO/Ctrl) indicates the % decrease necessary to account for the effect of the respective condition on MMM-*k*_HbO2_.

**Figure 11.**
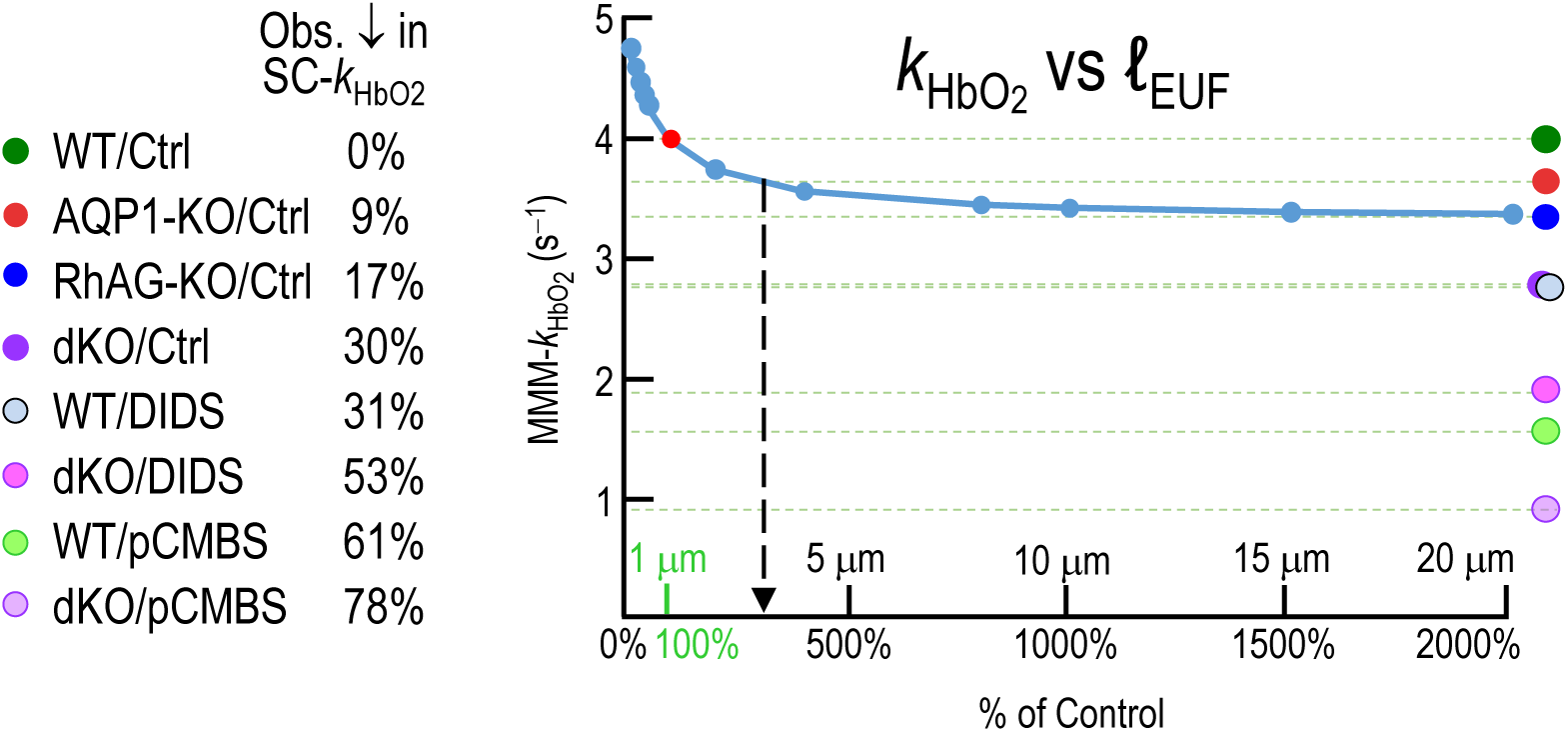
Mathematical simulations exploring the predicted sensitivity of *k*_HbO2_ to changes in thickness of the EUF Using our macroscopic mathematical model (MMM), we systematically varied the thickness of the EUF, ℓ_EUF_, to compute the corresponding rate constant for deoxygenation of HbO_2_ in intact RBCs (*k*_HbO2_). We computed the MMM-*k*_HbO2_ corresponding to the red dot—which corresponds to 100% of our standard ℓ_EUF_—by using the parameter values that come from Table 1. This is our standard condition for RBCs from wild-type mice in the absence of drugs (WT/Ctrl), and also is represented by the dark green circle and associated horizontal dashed green line. We performed eleven additional simulations (blue dots) by decreasing ℓ_EUF_ by the indicated percentages, keeping all other parameter values constant. The seven other colored circles and associated horizontal dashed lines indicate the MMM-*k*_HbO2_ values, relative to WT/Ctrl, for each of the other 7 experimental conditions. As summarized in the legend on the left, for each condition, we multiplied the standard MMM-*k*_HbO2_ (i.e., 3.99 s^−1^) by the shape-corrected decrease in the experimentally observed *k*_HbO2_ (e.g., ∼9% for AQP1-KO/Ctrl cells) to obtain the y-axis coordinate for that condition (e.g., 3.64 s^−1^ for AQP1-KO/Ctrl). These % decreases are also listed in the last row of Table 3. The downward arrow (tail on AQP1-KO/Ctrl) indicates the % decrease necessary to account for the effect of this condition on MMM-*k*_HbO2_.

**Figure 12.**
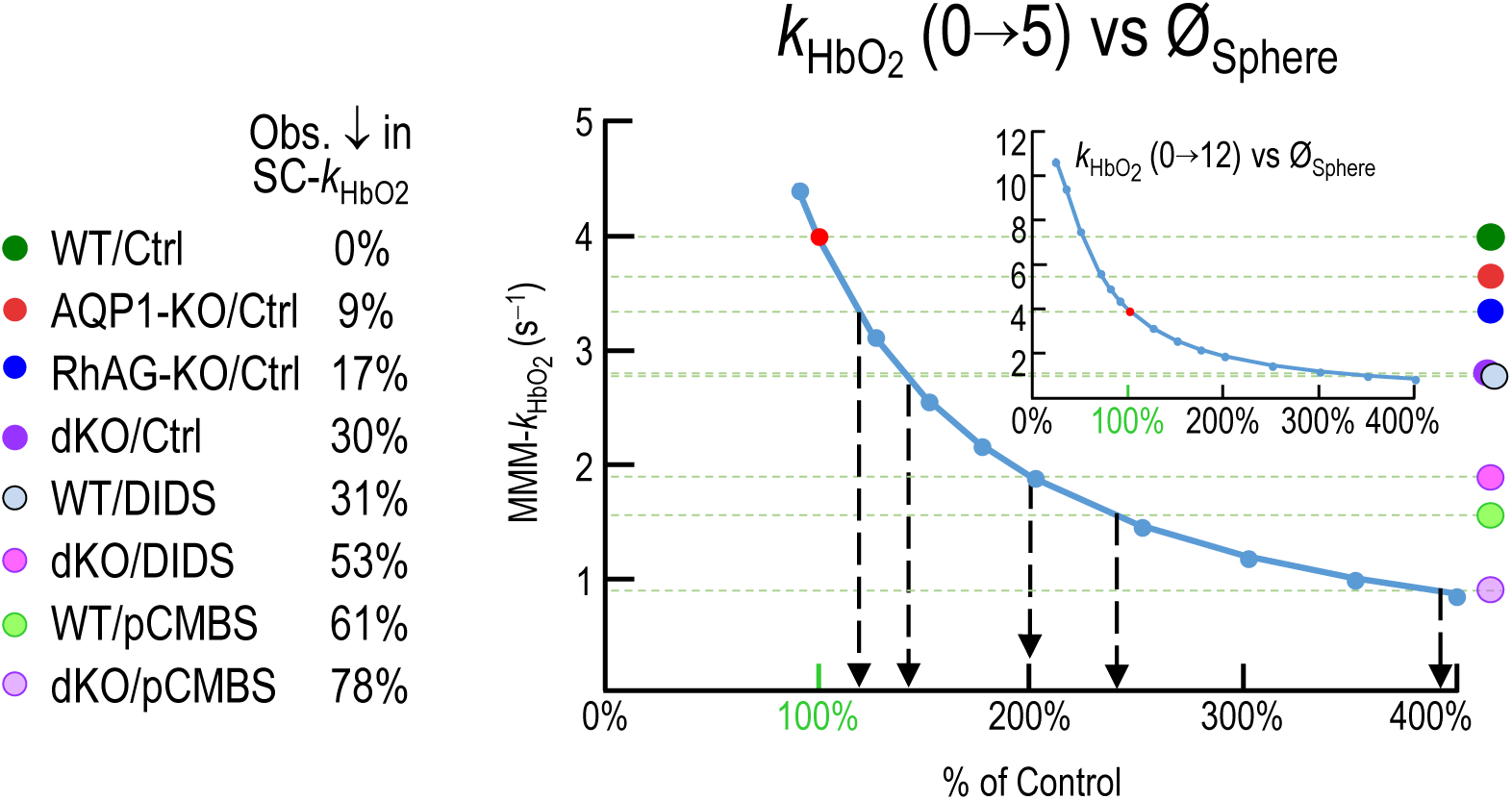
Mathematical simulations exploring the predicted sensitivity of *k*_HbO2_ to changes in sphere diameter Using our macroscopic mathematical model (MMM), we systematically varied the diameter of the sphere, Ø_Sphere_, to compute the corresponding rate constant for deoxygenation of HbO_2_ in intact RBCs (*k*_HbO2_). We computed the MMM-*k*_HbO2_ corresponding to the red dot—which corresponds to 100% of our standard Ø_Sphere_—by using the parameter values that come from Table 1. This is our standard condition for RBCs from wild-type mice in the absence of drugs (WT/Ctrl), and also is represented by the dark green circle and associated horizontal dashed green line. We performed additional simulations (blue dots) by decreasing Ø_Sphere_ by the indicated percentages, keeping all other parameter values constant. The seven other colored circles and associated horizontal dashed lines indicate the MMM-*k*_HbO2_ values, relative to WT/Ctrl, for each of the other 7 experimental conditions. As summarized in the legend on the left, for each condition, we multiplied the standard MMM-*k*_HbO2_ (i.e., 3.99 s^−1^) by the shape-corrected decrease in the experimentally observed *k*_HbO2_(e.g., ∼9% for AQP1-KO/Ctrl cells) to obtain the y-axis coordinate for that condition (e.g., 3.64 s^−1^ for AQP1-KO/Ctrl). These % decreases are also listed in the last row of Table 3. Note that main panel shows the y-axis from 0 to 5 s^−1^, whereas the inset expands the y-axis to extend from 0 to 12 s^−1^. The downward arrows (tails on AQP1-KO/Ctrl, etc.) indicate the % decreases necessary to account for the effect of the respective condition on MMM-*k*_HbO2_.

In each of these figures or panels, we plot MMM-*k*_HbO2_ on the y-axis and the varied parameter (as a % of the standard parameter value) on the x-axis. Thus, when the standard value of the parameter = 100%, MMM-*k*_HbO2_ = 3.99 s^−1^. As noted in our introduction of Figure 5, here we assume that all RBCs are intact BCDs, as for SC-*k*_HbO2_ values in paper #1.

In each figure/panel, we present eight dashed horizontal lines, one for each of the experimental conditions, each coded with a colored circle with same meaning as in Figure 6. For the y-axis value of a horizontal dashed line, we scale the hemolysis-corrected MMM-*k*_HbO2_ of 3.99 s^−1^ for the WT/Ctrl by the experimental condition’s % decrease in SC-*k*_HbO2_ vs. WT/Ctrl (Table 3, bottom line). Thus, MMM-*k*_HbO2_ = 3.99 s^−1^ for WT/Ctrl (0% decrease) and falls to ∼0.9 s^−1^ for dKO/pCMBS (78% decrease in MMM-*k*_HbO2_).

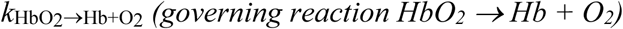

Figure 8 shows the dependence of MMM-*k*_HbO2_ on *k*_HbO2→Hb+O2_. As *k*_HbO2→Hb+O2_ rises, MMM-*k*_HbO2_ rises at first rapidly and then more slowly. The red dot represents the provisional control condition (i.e., WT/Ctrl) in which *k*_HbO2→Hb+O2_ is 100% of the standard parameter value (labeled in green on the x-axis) and MMM-*k*_HbO2_ is the familiar hemolysis-corrected value of 3.99 s^−1^. The condition with our smallest-observed % decrease in SC-*k*_HbO2_ is AQP1-KO/Ctrl (i.e., ∼9% lower than WT/Ctrl; Table 3). In order for a decrease in *k*_HbO2→Hb+O2_, by itself, to account for the observed ∼9% decrease in *k*_HbO2_, *k*_HbO2→Hb+O2_ would have to fall by ∼27% (i.e., x ≅ 73%; right downward dashed arrow). Our largest-observed % decrease in SC-*k*_HbO2_ was ∼78%, for dKO/pCMBS (see Table 3). In order for a decrease in *k*_HbO2→Hb+O2_, by itself, to lower the observed SC-*k*_HbO2_ by ∼78% (violet point and its horizontal dashed line), *k*_HbO2→Hb+O2_ would have to have fallen by ∼92% (i.e., x ≅ 8%; left downward dashed arrow).

In fact, our largest-observed decrease in *k*_HbO2→Hb+O2_ was ∼10% (see Table 2), for dKO/Ctrl. We conclude that the sensitivity of MMM-*k*_HbO2_ to decreases in *k*_HbO2→Hb+O2_ is far too small have an appreciable impact on our O_2_-offloading data.

### [Hb_Total_]_i_

Figure 9 shows the dependence of MMM-*k*_HbO2_ on [Hb_Total_]_i_. Starting from our lowest plotted value of x = 50% of the standard parameter value, increases in [Hb_Total_]_i_ cause MMM-*k*_HbO2_ to fall at first steeply and then more shallowly. The reasons for this slowing of O_2_ offloading are two-fold: (1) oxygen diffuses through the RBC far more slowly when bound to the large HbO_2_ tetramer than when free as O_2_ and (2) as [Hb_Total_]_i_ increases, total O_2_ increases—although [O_2_]_i_ does not—meaning that more time is required to rid the cell of O_2_.

In order to account for even our smallest-observed decrease in SC-*k*_HbO2_—that is, 9% (Table 3) in AQP1-KO/Ctrl—[Hb_Total_]_i_ would have to increase by 17% (i.e., x ≅ 117%; downward dashed arrow). In fact, in our genotype-specific comparisons (see Table 2), MCHC (analogous to [Hb_Total_]_i_) tended to fall in the KOs, with the greatest decrease being ∼5% in the AQP1-KOs/Ctrl. Thus, by themselves these decreases in MCHC would have caused an increase, rather than the observed decrease in SC-*k*_HbO2_. However, as noted in <u>paper #2</u>^33^, the decreases in MCHC were accompanied by increases in MCV, which have the opposite effect on SC-*k*_HbO2_ (as we will see <u>below</u>^27^). Thus, the MCHC and MCV effects tend to cancel.

In our drug studies (see <u>paper #2</u>^34^), pCMBS in dKO cells produced a 2% decrease—again in the wrong direction to explain a decrease in SC-*k*_HbO2_. On the other hand, pCMBS in WT cells produced a 4.4% increase in MCHC. However, examination of Figure 9 shows that increasing— [Hb_Total_]_i_ by 4.4% (i.e., x = 104.4%) would have only a trivial effect on MMM-*k*_HbO2_.

### Diffusion constants

Figure 10 shows the dependence of MMM-*k*_HbO2_ on the four relevant diffusion constants, namely, *D* for extracellular O_2_ (the only relevant extracellular solute), and the three intracellular diffusion constants for the components of the equilibrium HbO_2_ ^⇒^ Hb + O_2_.

### Extracellular *D*O_2_

Figure 10A shows that, starting from our standard parameter value for extracellular *D*_O2_ (i.e., 1.3313 × 10^−5^ cm^2^ s^−1^; see Table 1)—indicated by the red dot—decreasing (*D*_O2_)_o_ at first has little effect, and then causes MMM-*k*_HbO2_ to fall steeply. However, (*D*_O2_)_o_— which almost certainly does not vary substantially with genotype—would have to fall by ∼34% (i.e., x ≅ 66%; right downward dashed arrow) to account for the fall in SC-*k*_HbO2_ in our AQP1-KO/Ctrl experiments, and by ∼94% (i.e., x ≅ 6%; left downward dashed arrow) to account for the dKO/pCMBS data.

Because (*D*_O2_)_o_ governs the ability of O_2_ to diffuse through the EUF, away from the membrane surface to the bECF—where [O_2_] = 0 to mimic the annihilation of O_2_ by sodium dithionite or (Na^+^)_2_S_2_O^=^ (NDT)—we are not surprised that MMM-*k*_HbO4_ approaches zero as (*D*_O2_)_o2_ approaches zero. Thus, if (*D*_O2_)_o_ were zero, O_2_ offloading would stop as soon as [O_2_] at the outer surface of the membrane rose to match [O_2_]_i_.

Note that our model does not explicitly take into account the diffusion of NDT from the bECF into the EUF. Although the diffusion constant for S_2_O^=^ is doubtlessly lower than that for O_2_, is ∼71-fold greater than the initial [O_2_]_i_. Thus, we expect the EUF to be flooded with S_2_O^=^—to annihilate O_2_ as it exits the RBC) very early in the physiological experiment. It is for this reason that others believe that ℓ_EUF_ is probably <1 μm (Huxley & Kutchai, 1981; Vandegriff & Olson, 1984*b*; Holland *et al*., 1985), which, if true, means that (*D*_O2_)_o_ is even less impactful than indicated in Figure 10*A*.

### Intracellular *D*O_2_

Figure 10B shows that the dependence of MMM-*k*_HbO2_ on (*D*_O2_)_i_ is similar to that of (*D*_O2_)_o_ in Figure 10*A*. In fact, starting from the red dot in Figure 10*B*, which represents our standard parameter value for intracellular *D*_O2_ (i.e., 2.7745 × 10^−6^ cm^2^ s^−1^; see Table 1), decreasing (*D*_O2_)_i_ causes MMM-*k*_HbO2_ to fall somewhat less steeply than does decreasing (*D*_O2_)_o_ in panel *A*. Thus, (*D*_O2_)_i_—which again we regard as unlikely to change substantially with genotype— would need to fall by ∼40% (i.e., x ≅ 60%; right downward dashed arrow) to account for the fall in SC-*k*_HbO2_ in our AQP1-KO/Ctrl experiments, and by ∼98% (i.e., x ≅ 2%; left downward dashed arrow) to account for the dKO/pCMBS data.

Note that, as noted in <u>Methods</u>^35^, our choice of (*D*_O2_)_i_ is already, historically, a rather low value. If (*D*_O2_)_i_ is in fact greater than we estimated, then the red dot would slide further to the right (i.e., on an even flatter part of the curve, then the MMM-*k*_HbO2_ would be even less sensitive to changes in (*D*_O2_)_i_.

### Intracellular *D*HbO_2_

Figure 10C shows that MMM-*k*_HbO2_ is nearly insensitive to changes in *D*_HbO2_ until, starting at the red dot, this parameter falls to a tiny fraction of its standard parameter value (i.e., 6.07 × 10^−8^ cm^2^ s^−1^ cm s^−1^; see Table 1). The reason is that the diffusion constant of tetrameric HbO_2_ is so low that—even though the concentration of HbO_2_ is high—the movement of HbO_2_ per se toward the membrane during O_2_ offloading has little effect on O_2_ delivery, and thus to MMM-*k*_HbO2_. Of course, in Figure 9 we saw that decreases in [Hb_Total_]_i_ can greatly increase MMM-*k*_HbO2_. The reason is that Hb acts as a near-stationary buffer that binds O_2_ and nearly immobilizes it until O_2_ release from HbO_2_ permits the now-free O_2_ to resume its net diffusion to the membrane or to the next unoccupied Hb. Thus, we expect genotype-dependent changes in *D*_HbO2_ to have only miniscule effects on SC-*k*_HbO2_ until *D*_HbO2_ becomes so small that HbO_2_ approaches absolute immobility.

### Intracellular *D*Hb

Figure 10D shows that MMM-*k*_HbO2_ is even less sensitive to changes in *D*_Hb_ than to *D*_HbO2_. The reasons are: (1) The diffusion constant for the deoxygenated Hb tetramer (i.e., standard parameter value = 6.07 × 10^−8^ cm^2^ s^−1^; see Table 1) is nearly as low as it is for the fully oxygenated tetramer. And (2) Hb is moving away from the membrane during O_2_ offloading.

Thus, even if *D*_Hb_ were zero, the empty tetramers would pile up—at least mathematically—at the inner surface of the membrane, still maintaining a gradient for HbO_2_ tetramers to diffuse toward the membrane. At *D*_Hb_ = 0, the small decrease in MMM-*k*_HbO2_ would reflect absent diffusion of free Hb into the depth of the cytosol, and thus decreased formation of new HbO_2_.

### *ℓ*_EUF_

Figure 11 shows that MMM-*k*_HbO2_ has only weak dependence on ℓ_EUF_. Starting at x = 100% of the standard parameter value (i.e., ℓ_EUF_ = 1 μm), decreasing ℓ_EUF_ to as low as 90% of control (i.e., 0.1 μm) could raise SC-*k*_HbO2_ by <25%. Could an increase in ℓ_EUF_ account for our physiological data? By itself, a ∼3× increase in ℓ_EUF_ would be needed to replicate the SC-*k*_HbO2_ decrease that we see with AQP1-KO/Ctrl (see downward dashed arrow), and a ∼20× increase would be needed to replicate the RhAG-KO. No amount of ℓ_EUF_ increase, alone, could account for our other physiological data.

### *Ø*_Sphere_

#### Theoretical dependence of *k*HbO2 on Øsphere

The main part of Figure 12 and its inset (with a greatly expanded y-axis) show that MMM-*k*_HbO2_ is rather sensitive to changes in the diameter of the equivalent sphere because of the impact of O_2_ diffusion distance. The tails of the downward arrows are at the intersections of the blue MMM-*k*_HbO2_ curve and their respective observed SC-*k*_HbO2_ values (i.e., horizontal dashed lines):

#### Arrow #1

Our largest Ø_Sphere_ was for RhAG-KO/Ctrl, namely 2.18 μm (Table 2, data row 5), which is ∼7.9% greater than the value of 2.02 μm for WT/Ctrl. According to the curve in Figure ^12^ (see leftmost downward arrow), the percent increase in Ø_Sphere_ would have had to have been nearly 3× greater—about 20%—to account, by itself, for the observed decrease in SC-*k*_HbO2_.

#### Arrow #2

Our second largest computed Ø_Sphere_ was for dKO/Ctrl, namely 2.14 μm (Table 2, data row 5), which is only ∼5.9% greater than the WT/Ctrl value. According to our sensitivity analysis, the percent increase in Ø_Sphere_ would have to have risen by nearly 7×—by somewhat more than 40% (i.e., x ≅ 142%; 2^nd^ downward arrow)—in order to account for the observed decrease

Recall that the analyses in Figure 5—which take into account not just Ø_Sphere_ but all genotype-specific data—predict only a ∼4% decrease for RhAG-KO/Ctrl (vs. observed ∼17%), and a ∼4% to ∼6% decrease for dKO/Ctrl (vs. observed ∼30%).

#### Arrows #3, #4, #5

At greater increases in Ø_Sphere_, the curve flattens, so that one would have to invoke a ∼100% increase (i.e., x ≅ 200%; 3^rd^ downward arrow) to account for the dKO/DIDS data, more than a 140% increase (i.e., x > 240%; 4^th^ downward arrow) to account for the WT/pCMBS data, and nearly a 300% increase (i.e., x > 394%; rightmost downward arrow) to account for the dKO/pCMBS results. All of these values are unrealistic. We conclude that, by themselves, changes in Ø_Sphere_ cannot account for our physiological data.

### Tendency of MCHC and MCV effects to cancel

As already noted, in RBCs from KO strains, MCHC (a surrogate for [Hb_Total_]_i_; see <u>above</u>^36^) tends to fall, thereby speeding O_2_ offloading. On the other hand, as we just saw here, MCV (related to Ø_Sphere_) tends to rise, thereby slowing O_2_ offloading. These two effects tend to cancel, which is a reason that we see relative stability among the predicted MMM-*k*_HbO2_ values among genotypes in Figure 5.

### Accommodation for non-BCDs

The inputs to the MMM simulation include four observed experimental parameters: (1) MCV and (2) MCH (from which we compute MCHC), both of which come from automated hematology (<u>paper #1</u>^37^); (3) Ø_Major_, from imaging flow cytometry (<u>paper #2</u>^38^); and (4) *k*_HbO2→Hb+O2_, from stopped-flow absorbance spectroscopy on lysed RBCs (<u>paper #1</u>^39^). These values reflect all cells in the blood sample. An additional input is an estimate of *P*_M,O2_. The output of the simulation is *k*_HbO2_. Because the blood comprises both the dominant biconcave disks and minority non-BCDs, the paired values *P*_M,O2_ (input) and *k*_HbO2_ (output) represent hybrid values that to varying extents (depending on the sample) conflate BCDs and nBCDs. Our goal in this section is to obtain a (*P*_M,O2_, *k*_HbO2_) pair that estimates as accurately as possible the values for BCDs.

### Provisional estimates of P_M,O2_ and MMM-k_HbO2_ (steps #13 and #14 & 14′ in *Figure 1*)

Table 4 is an expanded version of Table 3. The first two data rows of Table 4 lists the prevalences of BCDs and nBCDs from <u>paper #2</u>^40^; we consider these in the next sub-section.

**Table 4.**
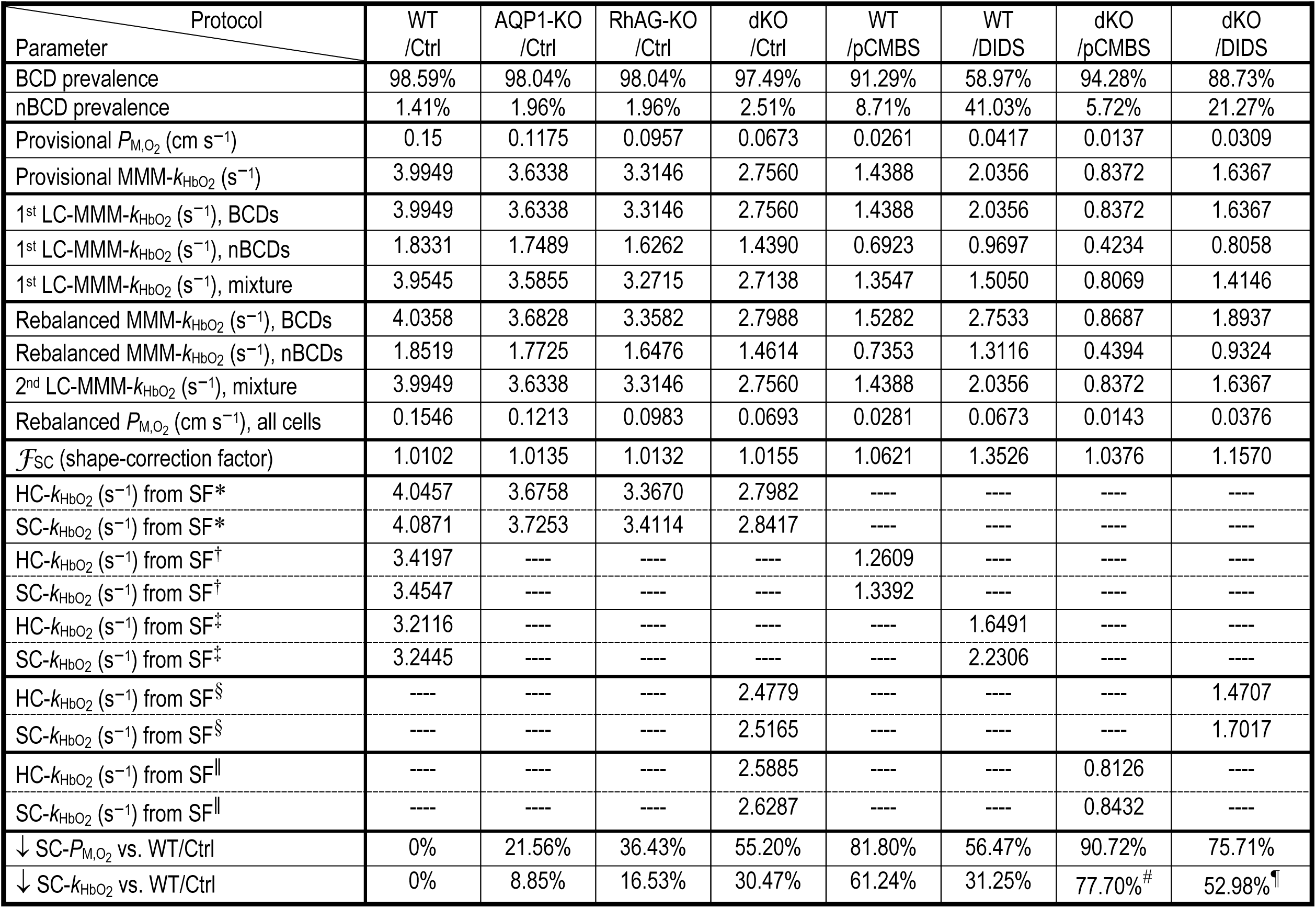
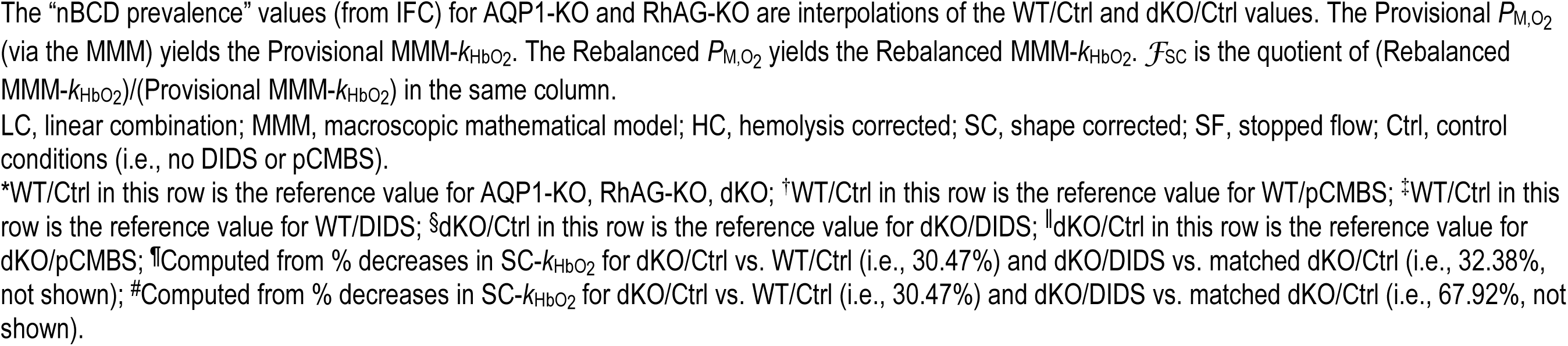
Detailed parameter values predicted by macroscopic mathematical modeling.

| Protocol<br>Parameter | WT<br>/Ctrl | AQP1-KO<br>/Ctrl | RhAG-KO<br>/Ctrl | dKO<br>/Ctrl | WT<br>/pCMBS | WT<br>/DIDS | dKO<br>/pCMBS | dKO<br>/DIDS |
| --- | --- | --- | --- | --- | --- | --- | --- | --- |
| BCD prevalence | 98.59% | 98.04% | 98.04% | 97.49% | 91.29% | 58.97% | 94.28% | 88.73% |
| nBCD prevalence | 1.41% | 1.96% | 1.96% | 2.51% | 8.71% | 41.03% | 5.72% | 21.27% |
| Provisional $P_{M,O_2}$ (cm s<sup>-1</sup>) | 0.15 | 0.1175 | 0.0957 | 0.0673 | 0.0261 | 0.0417 | 0.0137 | 0.0309 |
| Provisional MMM-$k_{HbO_2}$ (s<sup>-1</sup>) | 3.9949 | 3.6338 | 3.3146 | 2.7560 | 1.4388 | 2.0356 | 0.8372 | 1.6367 |
| 1<sup>st</sup> LC-MMM-$k_{HbO_2}$ (s<sup>-1</sup>), BCDs | 3.9949 | 3.6338 | 3.3146 | 2.7560 | 1.4388 | 2.0356 | 0.8372 | 1.6367 |
| 1<sup>st</sup> LC-MMM-$k_{HbO_2}$ (s<sup>-1</sup>), nBCDs | 1.8331 | 1.7489 | 1.6262 | 1.4390 | 0.6923 | 0.9697 | 0.4234 | 0.8058 |
| 1<sup>st</sup> LC-MMM-$k_{HbO_2}$ (s<sup>-1</sup>), mixture | 3.9545 | 3.5855 | 3.2715 | 2.7138 | 1.3547 | 1.5050 | 0.8069 | 1.4146 |
| Rebalanced MMM-$k_{HbO_2}$ (s<sup>-1</sup>), BCDs | 4.0358 | 3.6828 | 3.3582 | 2.7988 | 1.5282 | 2.7533 | 0.8687 | 1.8937 |
| Rebalanced MMM-$k_{HbO_2}$ (s<sup>-1</sup>), nBCDs | 1.8519 | 1.7725 | 1.6476 | 1.4614 | 0.7353 | 1.3116 | 0.4394 | 0.9324 |
| 2<sup>nd</sup> LC-MMM-$k_{HbO_2}$ (s<sup>-1</sup>), mixture | 3.9949 | 3.6338 | 3.3146 | 2.7560 | 1.4388 | 2.0356 | 0.8372 | 1.6367 |
| Rebalanced $P_{M,O_2}$ (cm s<sup>-1</sup>), all cells | 0.1546 | 0.1213 | 0.0983 | 0.0693 | 0.0281 | 0.0673 | 0.0143 | 0.0376 |
| $\mathcal{F}_{SC}$ (shape-correction factor) | 1.0102 | 1.0135 | 1.0132 | 1.0155 | 1.0621 | 1.3526 | 1.0376 | 1.1570 |
| HC-$k_{HbO_2}$ (s<sup>-1</sup>) from SF* | 4.0457 | 3.6758 | 3.3670 | 2.7982 | ---- | ---- | ---- | ---- |
| SC-$k_{HbO_2}$ (s<sup>-1</sup>) from SF* | 4.0871 | 3.7253 | 3.4114 | 2.8417 | ---- | ---- | ---- | ---- |
| HC-$k_{HbO_2}$ (s<sup>-1</sup>) from SF<sup>†</sup> | 3.4197 | ---- | ---- | ---- | 1.2609 | ---- | ---- | ---- |
| SC-$k_{HbO_2}$ (s<sup>-1</sup>) from SF<sup>†</sup> | 3.4547 | ---- | ---- | ---- | 1.3392 | ---- | ---- | ---- |
| HC-$k_{HbO_2}$ (s<sup>-1</sup>) from SF<sup>‡</sup> | 3.2116 | ---- | ---- | ---- | ---- | 1.6491 | ---- | ---- |
| SC-$k_{HbO_2}$ (s<sup>-1</sup>) from SF<sup>‡</sup> | 3.2445 | ---- | ---- | ---- | ---- | 2.2306 | ---- | ---- |
| HC-$k_{HbO_2}$ (s<sup>-1</sup>) from SF<sup>§</sup> | ---- | ---- | ---- | 2.4779 | ---- | ---- | ---- | 1.4707 |
| SC-$k_{HbO_2}$ (s<sup>-1</sup>) from SF<sup>§</sup> | ---- | ---- | ---- | 2.5165 | ---- | ---- | ---- | 1.7017 |
| HC- $k_{HbO_2}$ (s<sup>-1</sup>) from SF | ---- | ---- | ---- | 2.5885 | ---- | ---- | 0.8126 | ---- |
| SC- $k_{HbO_2}$ (s<sup>-1</sup>) from SF | ---- | ---- | ---- | 2.6287 | ---- | ---- | 0.8432 | ---- |
| ↓ SC-$P_{M,O_2}$ vs. WT/Ctrl | 0% | 21.56% | 36.43% | 55.20% | 81.80% | 56.47% | 90.72% | 75.71% |
| ↓ SC-$k_{HbO_2}$ vs. WT/Ctrl | 0% | 8.85% | 16.53% | 30.47% | 61.24% | 31.25% | 77.70%<sup>#</sup> | 52.98%<sup>¶</sup> |

The “nBCD prevalence” values (from IFC) for AQP1-KO and RhAG-KO are interpolations of the WT/Ctrl and dKO/Ctrl values. The Provisional $P_{M,O_2}$ (via the MMM) yields the Provisional $MMM-k_{HbO_2}$ . The Rebalanced $P_{M,O_2}$ yields the Rebalanced $MMM-k_{HbO_2}$ . $\mathcal{F}_{SC}$ is the quotient of (Rebalanced $MMM-k_{HbO_2}$ )/(Provisional $MMM-k_{HbO_2}$ ) in the same column.
LC, linear combination; MMM, macroscopic mathematical model; HC, hemolysis corrected; SC, shape corrected; SF, stopped flow; Ctrl, control conditions (i.e., no DIDS or pCMBS).
\*WT/Ctrl in this row is the reference value for AQP1-KO, RhAG-KO, dKO; <sup>†</sup>WT/Ctrl in this row is the reference value for WT/pCMBS; <sup>‡</sup>WT/Ctrl in this row is the reference value for WT/DIDS; <sup>§</sup>dKO/Ctrl in this row is the reference value for dKO/DIDS; <sup>||</sup>dKO/Ctrl in this row is the reference value for dKO/pCMBS; <sup>¶</sup>Computed from % decreases in $SC-k_{HbO_2}$ for dKO/Ctrl vs. WT/Ctrl (i.e., 30.47%) and dKO/DIDS vs. matched dKO/Ctrl (i.e., 32.38%, not shown); <sup>#</sup>Computed from % decreases in $SC-k_{HbO_2}$ for dKO/Ctrl vs. WT/Ctrl (i.e., 30.47%) and dKO/DIDS vs. matched dKO/Ctrl (i.e., 67.92%, not shown).

As noted in Methods, we chose our provisional *P*_M,O2_ for WT/Ctrl cells (Table 4, data row 3) to match the *P*_M,CO2_ value of 0.15 cm s^−1^ that Endeward et al (Endeward *et al*., 2006) obtained for the membrane CO_2_ permeability of human RBCs. MMM simulations based on *P*_M,O2_ = 0.15 cm s^−1^ yield a provisional MMM-*k*_HbO2_ of 3.99 s^−1^ (row 4), a value that is rather close to the highest of the three shape-corrected stopped-flow values (i.e., ∼4.09 s^−1^; row 14 in Table 4) for three separate WT/Ctrl data sets that we employed in <u>paper #1</u>^41^.

For the seven columns to the right of “WT/Ctrl” in Table 4, we chose the provisional *P*_M,O2_ values in iterative processes in which each *P*_M,O2_ generated a corresponding provisional MMM-*k*_HbO2_ that approximately matched the experimentally determined HC-*k*_HbO2_ value. Note that all (*P*_M,O2_, *k*_HbO2_) pairs lie on the sigmoidal *k*_HbO2_ vs. log(*P*_M,O2_) curve in Figure 6.

### nBCD prevalence (steps #15 & 15′ in *Figure 1*)

Recall that the first two data rows of Table 4 lists the prevalences of BCDs and nBCDs. The values for WT/Ctrl and dKO/Ctrl are around 2%, and may overestimate the actual values in SF experiments because of the extra time needed to prepare the samples for IFC analyses (compared to SF experiments; discussed in <u>paper #2</u>^42^). The nBCD prevalence values for the AQP1-KO/Ctrl and RhAG-KO/Ctrl groups are interpolations of the WT/Ctrl and dKO/Ctrl values. The nBCD prevalence values are ∼6-fold higher for the WT/pCMBS cells than the corresponding WT/Ctrl cells. The dKO/pCMBS values are only ∼2-fold greater than the corresponding dKO/Ctrl cells, probably reflecting the partial absence of pCMBS targets after the elimination of AQP1 and Rh_Cx_. The largest nBCD prevalence comes with WT/DIDS cells, at ∼41%; the figure falls to ∼21% in the dKO/DIDS group. These large percentages probably reflect the long incubation with DIDS (1 hour), compounded—as noted above—by the extra time needed for executing the IFC studies.

In the next four sections, we present a novel, 2-step approach for deconvoluting the HC-*k*_HbO2_ data on a mixture of BCDs and nBCDs to extract an estimate of the *k*_HbO2_ of BCDs, and thereby arrive at a *P*_M,O2_ that we assume to apply to both BCDs and nBCDs.

### First linear-combination analysis of k_HbO2_ values (step #16 in *Figure 1*)

Using WT/Ctrl cells to illustrate our approach, we begin by making the (incorrect) assumption that the provisional MMM-*k*_HbO2_ of ∼3.99 s^−1^ (Table 4, row 4) represents only BCDs (note that row 5 repeats row 4). This allows us to construct an idealized time course of HbO_2_ desaturation for BCDs (Figure 13*A*, green curve/furthest to the left). In parallel, knowing the measured Ø_Major_ of nBCDs, and assuming them to be spheres with the same *P*_M,O2_ as the BCDs (i.e., 0.15 cm s^−1^), we construct another idealized desaturation time course, but this time for nBCDs (Figure 13*A*, blue curve/furthest to the right). We then linearly combine these two curves in proportion to the prevalence of their respective cells. Because the weighting is ∼98.59% BCD (rows 5: LC-MMM-*k*_HbO2_ = 3.99 s^−1^) and 1.41% nBCD (rows 6: LC-MMM-*k*_HbO2_ = 1.83 s^−1^), the curve (Figure 13*A*, black) that describes the mixture of BCDs+nBCDs (rows 7: LC-MMM-*k*_HbO2_ = 3.95 s^−1^) lies barely to the right of the green BCD curve.

**Figure 13.**
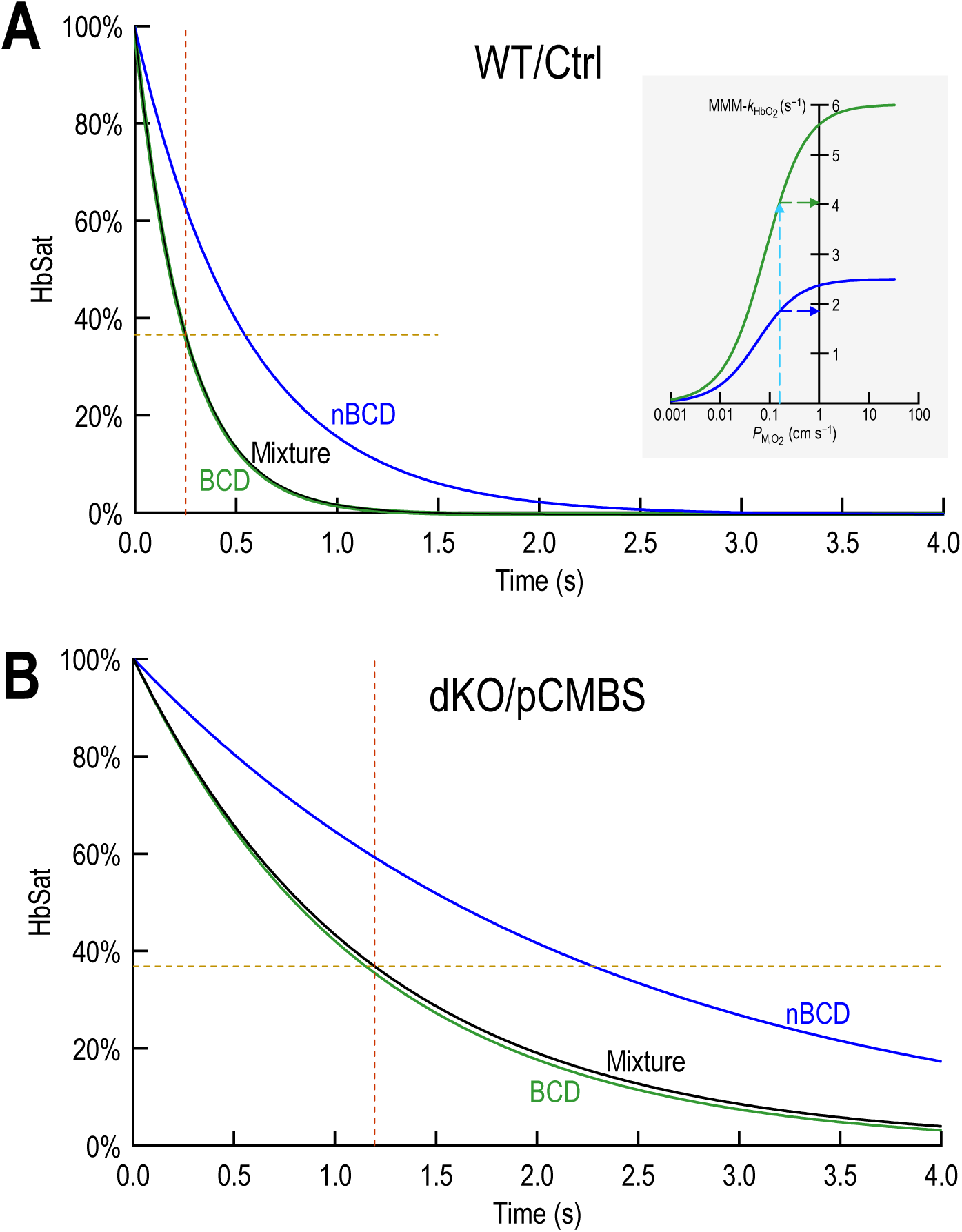
First linear-combination analysis of the rate constant for deoxygenation of HbO_2_, *k*_HbO2_ *A,* WT/Ctrl. The green curve is the computed decay of HbSat (hemoglobin saturation) for a biconcave disk (BCD). The blue curve is the computed decay for a non-BCD. The two sigmoid curves in the inset illustrate how, given a value for *P*_M,O2_, we compute the corresponding MMM-HbO_2_. The black curve, barely visible to the right of the green curve, is the linear combination of the green and blue curves, weighted for the respective prevalences of BCDs and nBCDs, respectively. The horizontal dashed curve represents the value of HbSat after 1 time constant for the mixture. The vertical dashed curve represents the corresponding time. *B,* dKO/pCMBS. Similar to panel *A* except here the calculations are for a dKO cell treated with pCMBS.

Figure 13B illustrates a similar analysis for dKO/pCMBS cells, at the opposite extreme of our experimentally observed spectrum of HC-*k*_HbO2_ values (Table 4, rows 13–22). Here, reflecting the low *P*_M,O2_, all three curves show a slower offloading of O_2_ from RBCs than in Figure 13*A*. Moreover, because the nBCD prevalence is ∼4-fold higher than for the WT/Ctrl cells in Figure 13*A*, the separation between the green (BCDs) and black (mixture) is more discernable.

### Rebalancing of k_HbO2_ values (steps #17 & 17′ in *Figure 1*)

Recall that in row 7 of Table 4, *k*_HbO2_ for the WT/Ctrl mixture is only ∼3.95 s^−1^, which is ∼1% lower than our provisional estimate (∼3.99 s^−1^), yet much higher than the *k*_HbO2_ for the nBCDs (1.83 s^−1^). The goal of the present step is to rebalance the three *k*_HbO2_ values so that the value for the mixture matches the original provisional *k*_HbO2_ as closely as possible. We make two assumptions:

- The term (MMM-*k*_HbO2/Mixture_ – MMM-*k*_HbO2/BCDs_)/(MMM-*k*_HbO2/BCDs_)) is maintained from the first linear combination throughout the rebalancing. That is, the rebalanced MMM-*k*_HbO2/BCDs_ exceeds the rebalanced MMM-*k*_HbO2/Mixture_ by the same fraction (∼1.02% for WT/Ctrl) as it did during the first linear-combination step.
- The ratio (MMM-*k*_HbO2/nBCDs_)/(MMM-*k*_HbO2/BCDs_) remains similarly fixed from the first linear combination through the rebalancing. For WT/Ctrl, this is ∼45.9%.

These provisos ensure that, after rebalancing, MMM-*k*_HbO2/Mixture_ is the same, within rounding errors, as our original provisional *k*_HbO2_, which, after all, really encompassed both BCDs and nBCDs. Thus, after rebalancing, MMM-*k*_HbO2/BCDs_ is slightly elevated (∼4.04 s^−1^ for WT/Ctrl). Achieving this higher *k*_HbO2_ requires that our corresponding rebalanced *P*_M,O2_ be increased from the provisional value—our next step.

### Rebalancing P_M,O2_ values (steps #18 – #20 in *Figure 1*)

Armed with the rebalanced MMM-*k*_HbO2_ (∼4.04 s^−1^ for WT/Ctrl in Table 4, row 8), we compute the rebalanced *P*_M,O2_ from an inverted/linearized version of Figure 6, shown in Figure 14. The result in the case of WT/Ctrl is a value (about “0.1546” cm s^−1^) that is slightly higher than our provisional *P*_M,O2_ of “0.1500” cm s^−1^.

**Figure 14.**
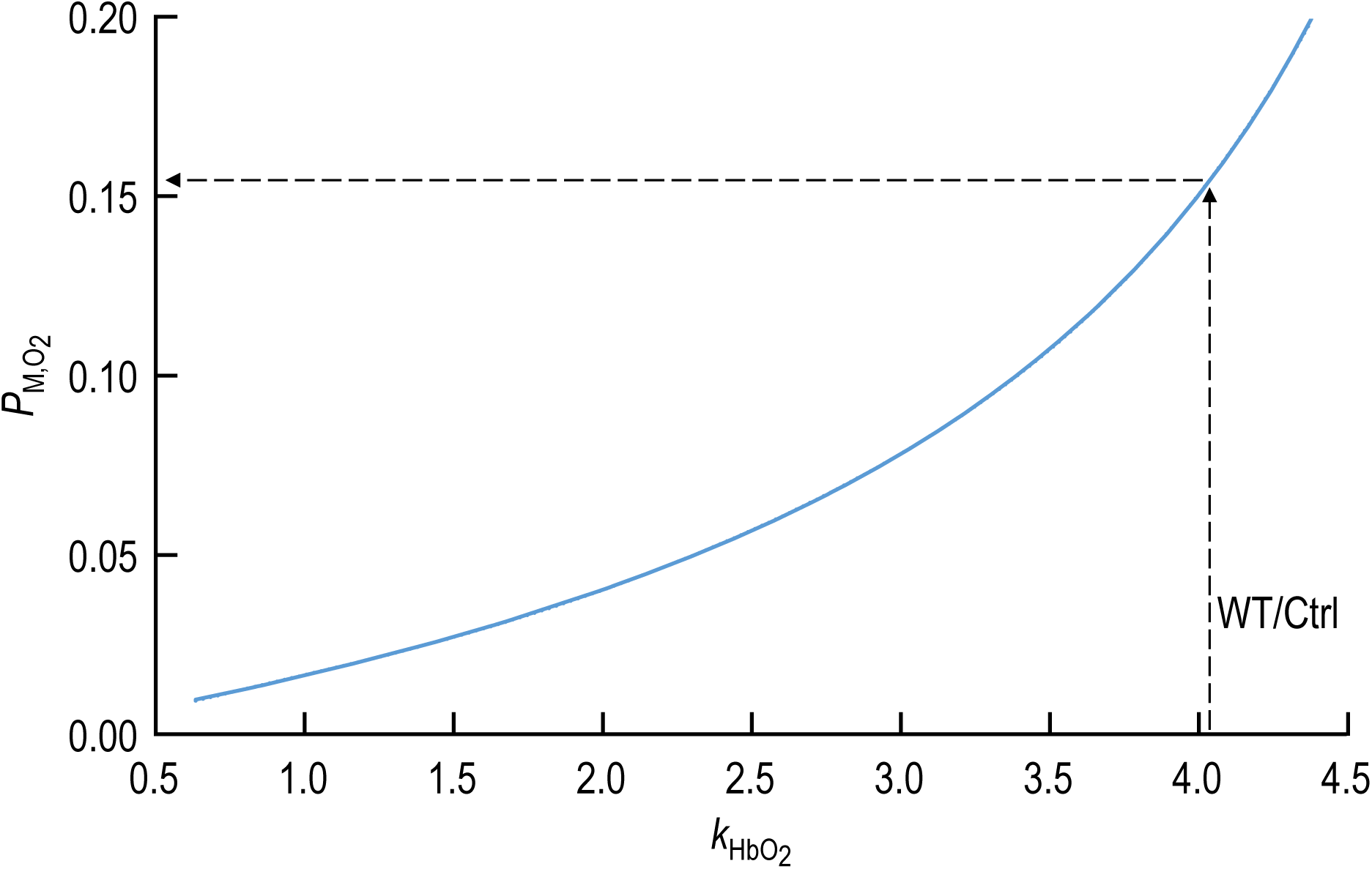
Inverted sigmoid plot This figure is analogous to Figure 5 except that here we plot *P*_M,O2_ on the y-axis, which is now linear, and the rate constant for deoxygenation of HbO_2_, *k*_HbO2_, on the x-axis. The upward-sloping curve is a best fit created from a 5^th^-order polynomial. This is a convenient tool for interpolating *P*_M,O2_ values that correspond to a particular *k*_HbO2_ value as part of the rebalancing of *P*_M,O2_. The upward arrow illustrates how, for a wild-type BCD under control conditions (no drugs), we identify the point on the curve that corresponds to the *P*_M,O2_ indicated by the leftward-pointing arrow.

### Shape-correction factor (F_SC_; steps #21 and #22 in *Figure 1*)

For a given data genotype/treatment, *F*_SC_ is the ratio (rebalanced MMM-*k*_HbO2_)/(provisional MMM-*k*_HbO2_), ∼1.0102 in the case of WT/Ctrl cells. Multiplying an HC-*k*_HbO2_ value (∼4.05 s^−1^ for the first set of WT/Ctrl data) by *F*_SC_ yields an estimate of SC-*k*_HbO2_ (∼4.09 s^−1^ in this example).

For dKO/pCMBS cells, with a higher nBCD prevalence than WT/Ctrl cells, *F*_SC_ is correspondingly higher, ∼1.0376.

The remainder of Table 4, following the *F*_SC_ values, shows five pairs of reference values (HC-*k*_HbO2_ and SC-*k*_HbO2_). The first set pertains to three data sets for control experiments: AQP1-KO/Ctrl, RhAG-KO/Ctrl, and dKO/Ctrl. The second set pertains to WT/pCMBS; the third, to WT/DIDS. The fourth and fifth sets are reference values for dKO/pCMBS and dKO/DIDS, respectively.

The last two rows of Table 4 show the percent decreases in HC-*k*_HbO2_ and SC-*k*_HbO2_, based on shape-corrected values, for each genotype/treatment, relative to WT/Ctrl. These are the same values listed above in the bottom two rows of Table 3.

## Discussion

To test whether the decreases in *k*_HbO2_ observed in paper #1 correspond to decreases in *P*_M,O2_ and, if so, to provide estimates of *P*_M,O2_ from measured *k*_HbO2_ values, we developed a three-dimensional reaction-diffusion model of O_2_ efflux from an RBC. In this model, we simplify the RBC by assuming that it is a perfectly symmetric sphere, the diameter of which equals the computed thickness of an RBC. Note that the thickness of the biconcave disk near its edge (∼2 μm)—not the major diameter (∼6.8 μm for mice)—is the dimension of the RBC (when it is a BCD) that is critically important for gas exchange, as pointed out by several authors (Roughton, 1932; Ponder, 1948; Forster, 1964; Endeward, 2012). Our approach, which has the advantage of geometric (and thus computational) simplicity, ignores the dimpled center of the BCD, which contains relatively little hemoglobin.

Our macroscopic mathematical model describes how the concentrations of O_2_, HbO_2_ and Hb change in time and space, when O_2_ diffuses out of the cytosol, across the plasma membrane, throughout the EUF and finally to the bECF. For our initial estimate of *P*_M,O2_ in our mathematical simulations, we assume that *P*_M,O2_ in mouse RBCs has the same value as measured by Endeward *et al*. (2006*b*) for the CO_2_ permeability of human RBCs.

Even though our MMM model is relatively simple and based only on first principles, our assumptions as well as the physiological data that inform the model are sufficiently robust that our model makes physiologically meaningful predictions for a WT mouse. One strength of this study is the use of the genotype-specific flow-cytometry and hematology data that we generated in the two accompanying papers to establish internally consistent parameter values.

### Krogh-Erlang Equation

In his pioneering modeling work on O_2_ delivery from capillary to tissue, August Krogh (1919) introduced a now-famous equation—derived by his mathematician-colleague Mr. K Erlang—that played a major role in elucidating the fundamental principles of how [O_2_] changes both longitudinally along the capillary and radially away from it. This equation reflects a highly simplified view of the system, which of course was necessary at that time, given the limitations in available computational algorithms and computing power. The analysis by Kreuzer (1982) lists 15 assumptions—some of which he and others had noted previously—that are implicit in the derivation of the Krogh-Erlang equation. Assumption #12 states that “the diffusion coefficient [of O_2_] is the same throughout the tissue.” Assumption #7 (previously noted by Hess, 1930)—really a corollary of #12—states that “the capillary wall does not present any resistance to oxygen diffusion.” Another corollary of #12 is that no cell membrane, including that of the RBC, offers resistance to O_2_ diffusion. We examine implications of this implicit assumption in the section <u>^b^elow</u>.

### How others concluded that the plasma membrane offers no resistance to O2 diffusion

After Krogh’s landmark paper, most practitioners in the RBC field adopted the aforementioned implicit assumption that membranes offer no resistance to O_2_ diffusion. However, a series of studies from Roughton and his colleagues led to the suggestion that the plasma membrane of RBCs offers significant resistance to the movement of gases, including CO_2_ and O_2_ (Blank & Roughton, 1960; Gibson *et al*., 1955; Forster, 1964; Roughton, 1964; Nicolson & Roughton, 1951; Lawson *et al*., 1965). However, later studies questioned this possibility and attributed the observed resistance to the extracellular unconvected fluid layer^43^ that surrounds the cells after flow stops in the SF apparatus (Kutchai, 1975; Coin & Olson, 1979; Huxley & Kutchai, 1981; Vandegriff & Olson, 1984*b*; Holland *et al*., 1985; Hughes & Bates, 2003). It is true that studies that focus on O_2_ influx across RBCs are intrinsically challenging because this influx depletes O_2_ at the surface of the RBC, and the EUF of that surrounds the RBC offers “resistance” to the replenishment of O_2_ from the bECF (Coin & Olson, 1979; Huxley & Kutchai, 1981; Holland *et al*., 1985; Hughes & Bates, 2003). Moreover, as pointed out by Vandegriff and Olson, the low solubility of O_2_ in water enhances the effect of EUFs in experiments of O_2_ influx (Vandegriff & Olson, 1984*b*). Nevertheless, it is possible that the implicit assumption of past authors—namely, that the plasma membrane offers no resistance of O_2_ diffusion (i.e., *R*_M,O2_—or *R*_M_ for short—is 0)—may have led them to dismiss Roughton hypothesis prematurely. The total resistance (*R*_Total_) to O_2_ diffusion from its original location in the cytosol to bECF is the sum of the effective resistance of the intracellular fluid (*R*_ICF_), *R*_M_, and the resistance of the extracellular unconvected fluid (*R*_EUF_). If one implicitly assumes that *R*_M_ ≅ 0, then the sum (*R*_M_ + *R*_EUF_) reduces to *R*_EUF_, with the result that one computes erroneously high values for the EUF thickness. In the process, one concludes that the large EUF explains the experimental data without the need of invoking membrane resistance—a classic bootstrap maneuver.

Because Vandegriff and Olson recognized the challenge of studying O_2_ influx, they investigated O_2_ efflux. They reduced the effects of the EUF by adding large amounts of sodium dithionite (NDT) in the suspending medium (Vandegriff & Olson, 1984*b*). The NDT near the outer side of the plasma membrane annihilates the O_2_ that has just left the RBC, thereby minimizing the build-up of O_2_ and minimizing the EUF. Viewed somewhat differently, the NDT maximizes the outwardly directed O_2_ gradient and therefore the rate of O_2_ efflux (Holland *et al*., 1985). Vandegriff and Olson used a reaction-diffusion model of an RBC, idealized as a cylindrical disk, to analyze the time course of their O_2_ release data. They concluded that their mathematical model—which lacked a term analogous to *R*_M_—could simulate their data without the need to include a membrane resistance to the movement of O_2_. Instead, they accounted for their O_2_-release data in the presence of NDT in the extracellular medium by including in the model an EUF, the width of which increased with time from the initial value of 1 μm to ∼30 μm (computed from an equation in ref. Holland *et al*., 1985). In summary, because they implicitly assumed *R*_M_ = 0, these authors invoked a very large *R*_EUF_, and used this result to conclude that *R*_M_ ≅ 0.

It is worth noting that at least 3 sets of authors have performed physicochemical experiments in which they concluded that lipid mono/bilayers can offer substantial resistance to CO_2_ or O_2_ (Blank & Roughton, 1960; Strutwolf *et al*., 2001; Ivanov *et al*., 2004).

### Recent observations on *D*O_2_ in the cytosol of RBCs

In their 2020 paper, Richardson et al conclude that the *D*_O2_ in the cytosol of RBCs is far lower than previously thought (Richardson *et al*., 2020). The generally accepted value for *D*_O2_ in RBC cytosol has been 5.09 × 10^−6^ cm^2^/ s^−1^ at 10 °C (Clark *et al*., 1985), which is ∼38% of the value in water (i.e., 13.313 × 10^-6^ cm^2^ s^−1^ at 10 °C). However, Richardson et al conclude that the actual *D*_O2_ value in RBC cytosol is <0.7 ×10^-6^ cm^2^ s^−1^ at 23 °C; we calculate that this value is < ∼3.5% of the value in water at 23 °C (i.e., 20 × 10^-6^ cm^2^ s^−1^; see Han & Bartels, 1996). Thus, the *D*_O2_ in cytoplasm predicted by Richardson et al should be 70/2000 = 1/29 of that in water.

### Our calculation of the *D*O_2_ of RBC cytosol (excluding membrane) at 10 °C

To apply the work of Richardson et al to our own, we must recompute their *D*_O2_ for a temperature of 10 °C. If we divide the aforementioned figure of 13.313 × 10^-6^ cm^2^ s^−1^ by 29, we arrive at a predicted value of *D*_O2_ (based on the work of Richardson et al) of 0.45907 × 10^−6^ cm^2^ s^−1^. We are concerned that this *D*_O2_ value is an underestimate because it is only ∼10% of the previously accepted value, a point recently made by Al-Samir *et al*. (2025). An inappropriately low estimate for (*D*_O2_)_i_ would impact us in two ways: (1) We would attribute to total diffusive resistance^44^

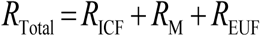

an ICF component that in fact is due to a combination of both the membrane and EUF layer. (2) Our standard *k*_HbO2_ value for WT/Ctrl would fall from 3.99 s^−1^—which comports very well with our physiological data—to only 2.61 s^−1^. Therefore, we decided to compromise by averaging the Richardson value of 0.45907 × 10^−6^ cm^2^ s^−1^ with the generally accepted value of 5.09 × 10^−6^ cm^2^ s^−1^ (Clark *et al*., 1985), and thereby arrive at a *D*_O2_ in RBC cytoplasm of 2.7745 × 10^−6^ cm^2^ s^−1^.

### Importance of values used in modeling

#### Thickness of EUF (ℓ_EUF_)

Several groups—Lawson *et al*. (1965), Coin and Olson (1979), Vandegriff and Olson (1984*b*), Yamaguchi *et al*. (1985), and Holland *et al*. (1985)—have emphasized the importance of employing a sufficiently high [NDT]_o_ to minimize the EUF layer in stopped-flow experiments. Lawson *et al*. (1965) concluded that the final NDT after mixing must be at least 10 mM, although this low value was later challenged on the basis of the “packet model” used in their data analysis. Coin and Olson (1979) showed that the “packet model” greatly overestimated the power of NDT, and concluded (see their figure 9) that an NDT of 25 mM is required to maximize *k*_HbO2_. Vandegriff and Olson (1984*b*) similarly discount the “packet model” and conclude that a final [NDT] of at least 25 mM is required. Yamaguchi *et al*. (1985), used a final [NDT] of 40 mM to maximize *k*_HbO2_.

In preliminary work, we systematically reduced [NDT] and noted a small *k*_HbO2_ falloff between [NDT]_Final_ values of 25 and 20 mM, and progressively steeper declines below 20 mM. Therefore, we chose a final [NDT] of 25 mM. We note that commercial supplies of NDT are not reliable, and we routine discard the majority of NDT bottles because of infiltration of air and H_2_O vapor (see <u>paper #1</u>^45^); NDT from such bottles routinely leads to lower *k*_HbO2_ values.

Despite the use of NDT in our SF experiments—which virtually abolishes the EUF (Holland *et al*., 1985)—in our mathematical model we make the conservative assumption that the EUF has a thickness of 1 μm and that no NDT is present in this layer. We simulate the annihilation of O_2_ by NDT by assuming that the bECF contains no O_2_ during the duration of the simulations. In Figure 11, we examine the dependence of *k*_HbO2_ on ℓ_EUF_. When we increase ℓ_EUF_, the simulated *k*_HbO2_ asymptotically falls to a value that approximates the biological value that we observe with RBCs from an RhAG-KO/Ctrl mouse. Thus, no increase in ℓ_EUF_ can account for our data with RBCs from dKO mice, or RBCs treated with pCMBS or DIDS.

Lowering ℓ_EUF_ to values smaller than 1 μm causes steep increases in the simulated *k*_HbO2_ (i.e., decreases in the resistance of the EUF to O_2_ diffusion, *R*_EUF_). Starting from such elevated *k*_HbO2_ values, the model can generate lower *k*_HbO2_ values (e.g., 3.99 s^−1^) by increasing *R*_M_, that is, by decreasing the permeability of the plasma membrane to O_2_ so that the sum (*R*_EUF_ + *R*_M_) remains the same. In other words, if the EUF is thinner than 1 μm—which is likely to be the case in the presence of high [NDT]—the membrane must make a greater contribution to the overall diffusive resistance of O_2_.

#### Calculation of the contribution of the plasma membrane to total diffusive resistance to O2

Let us assume that we can treat the diffusion of O_2_ from the ICF of the RBC to the bECF like an electrical circuit (see Figure 2*A*), where, as in the preceding equation, *R*_Total_ is the sum *R*_ICF_ + *R*_M_ + *R*_EUF_. In the next 3 sub-sections, we will compute each of the resistances on the right side of the equation, and then we will compute the fraction of *R*_Total_ that *R*_M_ represents. See Table 1 for parameter values used in simulations of RBCs from WT/Ctrl mice.

### Calculate *R*ICF

We start by computing—at the time *t*_37_ (i.e., 0.2503 s), when HbSat has fallen to 1/e of its initial value—the parallel fluxes of free O_2_ (*J*_O2_) and HbO_2_ (*J*_HbO2_) from the position of the average O_2_ and HbO_2_ molecules in the ICF. From these two fluxes, we then compute the total O_2_ flux (*J*_Total,O2_).

First, we determine the number of O_2_ molecules in each of the 101 shells (*i* = 1 to 101) that make up the model RBC, as well as the distance (ℓ) from the center of each shell to the inner surface of the plasma membrane (iM). For a WT cell with no inhibitors, we find that the average distance of an O_2_ molecule from the iM is 2.792×10^-5^ cm. For HbO_2_, a similar analysis yields an average distance of 2.560×10^-5^ cm. Because these distances are so similar, we used the average distance for O_2_ and HbO_2_, which is 2.68×10^-5^ cm, which corresponds to shell *i* = *k* = 76 from the center of the sphere. We call this average distance ℓ_k_. Thus, the flux of free O_2_ from ℓ_k_ to the iM is:

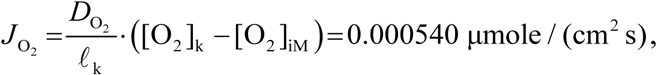

where *D*_O2_ is the diffusion constant for O_2_ in the ICF (see Table 1 for value), and where [O_2_]_k_ = 0.01528 mM and [O_2_]_iM_ = 0.01007 mM at *t*_37_. For HbO_2_,

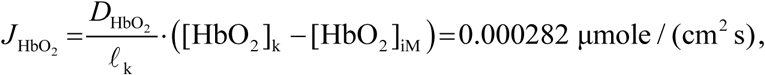

where *D*_HbO2_ is the diffusion constant for HbO_2_ in the ICF (see Table 1 for value), and where [HbO_2_]_k_ = 6.869 mM and [HbO_2_]_iM_ = 6.744 mM at *t*_37_.

Thus, the total O_2_ flux from ℓ_k_ to the iM is

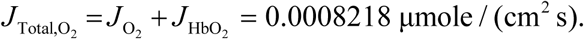

The effective resistance to the diffusion of O_2_ is^46^

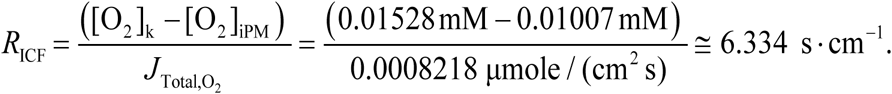

### Calculate *R*M

We calculate the resistance of the plasma membrane to O_2_ diffusion as the reciprocal of the “true” microscopic permeability of the plasma membrane to O_2_, which is 0.15 cm s^−1^ (see Table 1 for value). Thus,

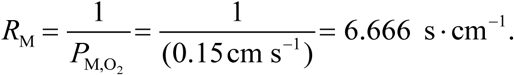

### Calculate *R*EUF

We calculate the resistance of the EUF to O_2_ diffusion as

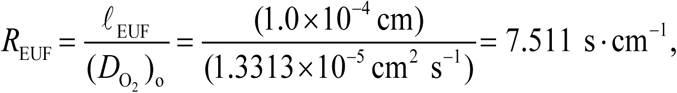

where ℓ_EUF_ is the thickness of the EUF and *D*_O2_ is the diffusion constant for O_2_ in the EUF (see Table 1 for values).

### Compute the contribution of *R*M to *R*Total

We calculate the contribution of *R*_M_ to *R*_Total_ as

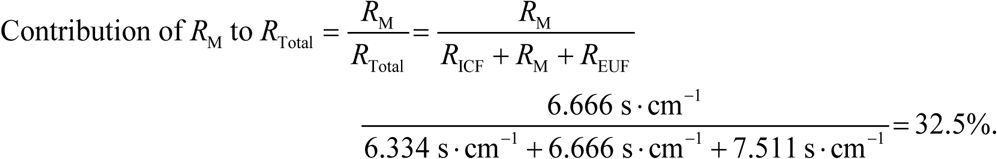

Note that if ℓ_EUF_ is in fact <1 μm (i.e., *R*_EUF_ < 7.511 s cm^−1^), the contribution of *R*_M_ to total resistance would be greater than ∼33%. Note also that if our compromise estimate for (*D*_O2_)_i_ is in fact too low (i.e., if we are overestimating *R*_ICF_), this consideration would also increase the contribution of *R*_M_ to *R*_Total_.

### Potential contribution of channels to *R*M and *R*total

In reaching the above conclusion that *R*_M_ represents at least ∼33% of *R*_Total_ for WT/Ctrl RBCs under the conditions of our experiments, we did not consider inhibitors or KO strains. Our observed 30% decrease in SC-*k*_HbO2_ caused by the dKO/Ctrl corresponds to a ∼55% decrease in *P*_M,O2_ (see Figure 6), so that *R*_M_ is now (6.666 s cm^−1^)/(1–0.55), which represents ∼52% of *R*_Total_. The 78% decrease in *k*_HbO2_ caused by dKO/pCMBS corresponds to a ∼91% decrease in *P*_M,O2_, so that *R*_M_ now rises to (6.666 s cm^−1^)/(0.55), so that *R*_M_ now is ∼84% of *R*_Total_—even with our conservative estimates of *R*_ICF_ (i.e., intracellular *D*_O2_) and *R*_EUF_ (i.e., ℓ_EUF_). Viewed differently, for WT/Ctrl RBCs under the conditions of our experiments, the reason that *R*_M_ makes only a ∼33% contribution to *R*_Total_ is that the O_2_ channels make the membrane leaky.

It is not clear how drugs confined to the outside of the membrane or the genetic deletion of membrane proteins could produce the observed decreases of *k*_HbO2_ in any way other than decreasing O_2_ egress through protein channels. Moreover, an analysis of 8 key parameters (e.g., ℓ_EUF_, cell thickness) based on mathematical simulations shows that no reasonable change in any could explain our *k*_HbO2_ data (Figure 8 through Figure 12).

### Uniqueness of the sigmoidal *k*HbO2 vs. log(*P*M,O_2_) curve

Our development of the sigmoid curve in Figure 6 is novel. The curve describes all possible combinations of *P*_M,O2_ and *k*_HbO2_ for a particular set of conditions. The detailed shape of the curve depends on experimental conditions (e.g., temperature) and MMM-simulation parameters (e.g., ℓ_EUF_) as summarized in Table 1. Beyond these values, the shape of the sigmoid curve is potentially unique for each species or strain of animal—or individual humans with unique genetics and medical histories—because each will have a characteristic set of genotype-specific values for MCV and MCH (and thus MCHC), Ø_Major_, and *k*_HbO2→Hb +O2_ (see Table 2). Thus, with different experimental conditions and strains/species, we expect to see major differences in the maximal-possible *k*_HbO2_ at the far right of the curve, which represents a membrane made of a film of water.

### Accommodation for nBCDs

Our accommodation for nBCDs is also novel, founded on our MMM-simulations with various algebraic manipulations, together with a meticulous set of collected data. For RBCs from our WT and KO strains—studied in the absence of drugs—the prevalence of nBCDs is small and thus the corrections are very minor. In the presence of pCMBS the corrections become larger, and for DIDS, still larger. The accuracy of the corrections likely increases as nBCD prevalence falls, and would improve with a more detailed knowledge of the surface-to-volume ratio.

## Acknowledgements

We thank Jean-Pierre Cartron for the gift of RhAG-KO mice. We thank him and Gerolf Gros and for extremely helpful discussions at the outset of the project. We also thank Pawel Swietach for comments on an earlier version of the manuscript. We thank Seong-Ki Lee for developing an improved approach for genotyping the RhAG-KO mice. We thank systems analyst Dale Huffman for using MATLAB to generate some of the figures. The authors gratefully acknowledge Daniela Calvetti and Erkki Somersalo for having developed the engine of an earlier version of the CO_2_/pH reaction-diffusion model of an oocyte, which in part served as the starting point for the RBC model. We thank Thomas Radford for organizing the husbandry of the mouse colonies; Gerald Babcock for his role as laboratory manager; James W. Jacobberger and Philip G. Woost of the CWRU Flow Cytometry and Imaging Microscopy Core (FCIMC) for their assistance with flow cytometry; and Daniela Schlatzer of the CWRU Center for Proteomics and Bioinformatics for their assistance with mass spectrometry. This work was supported by Office of Naval Research (ONR) grant N00014-11-1-0889, N00014-14-1-0716, and N00014-15-1-2060; a Multidisciplinary University Research Initiative (MURI) grant N00014-16-1-2535 from the DoD, NIH grant multi-scale modeling grant 5U01GM111251 (to WFB), and NIH grant R01HL160857 (to WFB). R.O. and the modeling were supported in part by NIH grant K01-DK107787. W.F.B. gratefully acknowledges the support of the Myers/Scarpa endowed chair.

## Additional Information

### Competing interests

The authors declare that they have no competing interests.

### Author contributions

R.O., P.Z. & W.F.B. designed the study; R.O. developed and implemented the computational model, and performed all numerical simulations; R.O., P.Z. and W.F. B. analyzed data. R.O., P.Z. & W.F.B. prepared the figures; R.O., P.Z., F.J.M., & W.F.B. wrote the manuscript.

## Footnotes

1 As a shorthand, we will refer to the three papers as “paper #1”, “paper #2”, and “paper #3”.

2 This manuscript was first published as a preprint: Occhipinti R, Zhao P, Moss FJ & Boron WF (2025). Role of channels in the O₂ permeability of murine red blood cells. III. Mathematical modeling and simulations. bioRxiv. https://www.biorxiv.org/content/10.1101/2025.03.05.639964

3 When we use “Hb”—as is standard practice in the literature—without using a subscript designation “T” for tetramer or “M” for monomer, we mean the mass of hemoglobin (in grams), both deoxyhemoglobin (Hb) and oxyhemoglobin (HbO2). Thus, [HbTotal]i is the total intracellular Hb concentration—both Hb and HbO2—measured in g/dl.

4 See Methods>Mathematical modeling and simulations>Calculation of RBC thickness based on the geometry of a torus. The “>” symbol separates heading levels. It is understood that the reference is to a location in the present paper (i.e., paper #3) unless “See” is followed by “Paper #1” or “Paper #2”.

5 Not to be confused with *r*Torus, which we use below to describe the minor radius of a torus.

6 See Paper #1>Methods>Determination of rate constant *k*HbO2→Hb+O2 (workflow #11)

7 These include immature RBCs (i.e., nucleated cells, reticulocytes, and spherocytes) as determined by imaging flow cytometry in paper #2.

8 See Paper #2>table 3

9 See Paper #2>table 4

10 Here we use “Total” to mean all Hb tetramers, whether they bind O2 (i.e., HbTO2) or not (i.e., HbT). Thus, [HbT,Total] = [HbTO2] + [HbT].

11 These hematology data pertain to the subset of mice used in the imaging flow cytometry and MMM studies.

12 Here we use “Total” to mean all Hb monomers, whether they bind O2 (i.e., HbMO2) or not (i.e., Hbm). Thus, [HbM,Total] = [HbMO2] + [HbM].

13 This is the “permeability” computed on the basis of concentrations immediately adjacent to the two sides of the membrane (i.e., corresponding to *R*Sphere– and *R*Sphere+). Physiologists usually define permeability operationally, based on the respective concentrations in the bulk intra- and extracellular fluids (see Boron, 2010).

14 *P*M,S is the general description for membrane permeability to solute *S*. When *S* is O2, elsewhere in the three papers, we refer to the permeability as *P*M,O_2_.

15 See Paper #1>Results>Effect of pCMBS or DIDS on O2 offloading from RBCs>Stopped-flow Approach

16 See (a) Paper #1>Methods>Physiological Solutions … & … (b)..>Stopped-flow absorbance spectroscopy (workflow #1)

17 See Discussion>Recent observations on *D*O_2_ in the cytosol of RBC

18 See Paper #1>Methods>Calculation of raw-*k*HbO2 (workflow #3)

19 See Methods>Mathematical modeling and simulations>Model formulation

20 See Paper #1>Results>Effect of genetic deletions on O2 offloading from RBCs

21 See Paper #1>figure 3*d*, dark-green curve

22 See Methods>Mathematical modeling and simulations>Simulation of time course of HbO2 deoxygenation, and estimation of *k*HbO2

23 See Results>Accommodation for non-BCDs

24 See Paper #1>figure 5*b*

25 See Paper #2>figure 8*b–c*, “Ctrl” bars

26 See Paper #1>figure 5*b*

27 See Results>Mathematical simulations exploring the predicted sensitivity of *k*HbO2 to eight key kinetic and geometric parameters

28 See Discussion>Uniqueness of the sigmoidal *k*HbO2 vs. log(*P*M,O_2_) curve

29 See Paper #2>table 5

30 See Results>Accommodation for non-BCDs

31 See Paper #2>Results>Proteomics

32 See Paper #1>figure 5*d*, light purple bar

33 See Paper #2>table 4

34 See Paper #2>table 6

35 See Methods>Mathematical modeling and simulations>Parameter values

36 See Results>Mathematical simulations exploring the predicted sensitivity …>[HbTotal]i

37 See Paper #1>Results>Hematological and related parameters >Automated hematology

38 See Paper #2>Results>Morphometry>Imaging flow cytometry

39 See Paper #1>Results>Hematological and related parameters>Hb kinetics (*k*HbO2→Hb+O2)

40 See Paper #2>table 3

41 See Paper #1>Results>Mathematical simulations>Summary

42 See Paper #2>Discussion>Morphometry>Imaging flow cytometry…>nBCD (poikilocyte) abundance >Time dependence

43 We avoid using the classic term “unstirred layer” because it is defined operationally; it has meaning only in the steady state, in principle differs for each solute, and is not easily implemented in MMM (see Figure 2*A*).

44 Note that this equation includes all diffusive terms but does not include the reaction term for the dissociation of HbO2

45 See Paper #1>Discussion>Implications of the macroscopic mathematical model>Basic features of the model>Simulated *k*HbO2 for WT/Ctrl RBCs

46 Apparent deviations in computed values are due to rounding errors

## Notes

### Competing Interest Statement

The authors have declared no competing interest.

### Summary of Updates

Minor revisions to Abstract text. Revised text of Results and Discussion New Figure 1 is added to demonstrate the workflow and interaction between the experiments and simulations between the 3 papers; the present paper #3 and associated paper #1 and paper #2. Paper #1: https://doi.org/10.1101/2025.03.05.639948 Paper #2: https://doi.org/10.1101/2025.03.05.639962 V1_Fig.1 becomes V2_Fig2 New Figure 3 to present the conversion of biconcave disk geometry into torus geometry. V1_Fig.1 becomes V2_Fig4 and now also displays two new panels. Panel C: Concentration of the sum of free O_2_ and HbO_2_ monomers, integrated over the entire volume of cytoplasm. Panel F: Transmembrane O_2_ flux (*J* _M,O2_). V1_Fig.3 becomes V2_Fig3 and has a new Panel B: displaying MMM-*k* _HbO2_ values predicted from simulations *k* _HbO2_ -->_Hb+O_2_ _ assumed to be specific for each genotype V1_Fig.4 becomes V2_Fig6 V1_Fig.5 becomes V2 Fig7: Fig7F now has an inset showing a magnification of the first 1 ms of the simulated J_M,O2_ V1_Fig.6 becomes V2_Fig8 and the symbol key is updated with revised nomenclature V1_Fig.7 becomes V2_Fig9 and the symbol key is updated with revised nomenclature V1_Fig.8 becomes V2_Fig10 and the symbol key is updated with revised nomenclature V1_Fig.9 becomes V2_Fig12 and the symbol key is updated with revised nomenclature V1_Fig.10 becomes V2_Fig12. The symbol key is updated with revised nomenclature, and the position of the inset is moved. V1_Fig.11 becomes V2_Fig13. Added annotations for a biconcave disk (BCD) and non-BCD for computed decay of HbSat (hemoglobin saturation). Also added annotation for the linear combination of the green and blue curves, "Mixture". V1_Fig.12 becomes V2_Fig14. The cross-references to Paper #1 and Paper #2, the two supporting preprints, are updated.

## References

Adair GS, Bock AV & Field H (1925). The hemoglobin system VI. the oxygen dissociation curve of hemoglobin. J Biol Chem 63, 529–545.

Al-Samir S, Kyriazi D, Yool AJ, Moser I, Kyriazi K, Gros G, Tsiavaliaris G & Endeward V (2025). Aquaporin-1 acts as an O₂ channel. The permeability of human and mouse red cell membranes for oxygen. Am J Physiol Cell Physiol; DOI: 10.1152/ajpcell.00858.2024.

Blank M & Roughton FJW (1960). The permeability of monolayers to carbon dioxide. Trans Faraday Soc 56, 1832–1841.

Boron WF (2010). Sharpey-Schafer lecture: Gas channels. Exp Physiol 95, 1107–1130.

Boron WF, Endeward V, Gros G, Musa-Aziz R & Pohl P (2011). Intrinsic CO₂ permeability of cell membranes and potential biological relevance of CO₂ channels. ChemPhysChem 12, 1017–1019.

Clark A, Federspiel WJ, Clark PA & Cokelet GR (1985). Oxygen delivery from red cells. Biophys J 47, 171–181.

Coin JT & Olson JS (1979). The rate of oxygen uptake by human red blood cells. J Biol Chem 254, 1178–1190.

Cooper GJ & Boron WF (1998). Effect of pCMBS on CO₂ permeability of *Xenopus* oocytes expressing aquaporin 1 or its C189S mutant. Am J Physiol 275, C1481–1486.

Cooper GJ, Occhipinti R & Boron WF (2015). CrossTalk proposal: Physiological CO₂ exchange can depend on membrane channels. J Physiol; DOI: 10.1113/JP270059.

Deonikar P, Abu-Soud HM & Kavdia M (2014). Computational analysis of nitric oxide biotransport to red blood cell in the presence of free hemoglobin and NO donor. Microvasc Res 95, 15–25.

Deonikar P & Kavdia M (2013). Contribution of membrane permeability and unstirred layer diffusion to nitric oxide-red blood cell interaction. J Theor Biol 317, 321–330.

Endeward V (2012). The rate of the deoxygenation reaction limits myoglobin- and hemoglobin-facilitated O₂ diffusion in cells. J Appl Physiol 112, 1466–1473.

Endeward V, Al-Samir S, Itel F & Gros G (2014). How does carbon dioxide permeate cell membranes? A discussion of concepts, results and methods. Front Physiol 4, 382.

Endeward V, Musa-Aziz R, Cooper GJ, Chen L-M, Pelletier MF, Virkki LV, Supuran CT, King LS, Boron WF & Gros G (2006). Evidence that aquaporin 1 is a major pathway for CO₂ transport across the human erythrocyte membrane. FASEB J 20, 1974–1981.

Fick A (1855). V. On liquid diffusion. Lond Edinb Dubl Phil Mag 10, 30–39.

Forster RE (1964). Rate of gas uptake by red cells. In Handbook of Physiology, pp. 827–837. American Physiological Society.

Forstner H & Gnaiger E (1983). Appendix A. Calculation of equilibrium oxygen concentration. In *Polarographic oxygen sensors: aquatic and physiological applications*. Springer-Verlag, Berlin Heidelberg New York.

Geers C & Gros G (2000). Carbon dioxide transport and carbonic anhydrase in blood and muscle. Physiol Rev 80, 681–715.

Gibson QH, Kreuzer F, Meda E & Roughton FJ (1955). The kinetics of human haemoglobin in solution and in the red cell at 37 degrees C. J Physiol (Lond) 129, 65–89.

Graham T (1829). Notice of the singular inflation of a bladder. Quart J Sci II, 88–89.

Graham T (1833). XXVII. On the law of the diffusion of gases. Lond Edinb Dubl Phil Mag 2, 175–190.

Han P & Bartels DM (1996). Temperature Dependence of Oxygen Diffusion in H₂O and D₂O. J Phys Chem 100, 5597–5602.

Hartridge H & Roughton FJ (1927). The rate of distribution of dissolved gases between the red blood corpuscle and its fluid environment: Part I. Preliminary experiments on the rate of uptake of oxygen and carbon monoxide by sheep’s corpuscles. J Physiol (Lond) 62, 232– 242.

Hess WR (1930). Die regulierung des Blutkreislaufes: gleichzeitig ein Beitrag zur Physiologie des vegetativen Nervensystemes. Thieme.

Holland RA, Shibata H, Scheid P & Piiper J (1985). Kinetics of O₂ uptake and release by red cells in stopped-flow apparatus: effects of unstirred layer. Respir Physiol 59, 71–91.

Huffman DE, Occhipinti R, Somersalo E, Calvetti D & Boron WF (2025). A graphical user interface for “Role of channels in the O2 permeability of murine red blood cells III. Mathematical modeling.” Available at: 10.5281/zenodo.14932553.

Hughes JMB & Bates DV (2003). Historical review: the carbon monoxide diffusing capacity (DLCO) and its membrane (DM) and red cell (Theta.Vc) components. Respir Physiol Neurobiol 138, 115–142.

Huxley VH & Kutchai H (1981). The effect of the red cell membrane and a diffusion boundary layer on the rate of oxygen uptake by human erythrocytes. J Physiol (Lond) 316, 75–83.

Ivanov II, Fedorov GE, Gus’kova RA, Ivanov KI & Rubin AB (2004). Permeability of lipid membranes to dioxygen. Biochem Biophys Res Commun 322, 746–750.

Kreuzer F (1982). Oxygen supply to tissues: the Krogh model and its assumptions. Experientia 38, 1415–1426.

Krogh A (1919). The number and distribution of capillaries in muscles with calculations of the oxygen pressure head necessary for supplying the tissue. J Physiol (Lond) 52, 409–415.

Kutchai H (1975). Role of the red cell membrane in oxygen uptake. Respir Physiol 23, 121–132.

Lawson WH, Holland RA & Forster RE (1965). Effect of temperature on deoxygenation rate of human red cells. J Appl Physiol 20, 912–918.

Michenkova M, Taki S, Blosser MC, Hwang HJ, Kowatz T, Moss FJ, Occhipinti R, Qin X, Sen S, Shinn E, Wang D-K, Zeise B, Zhao P, Malmstadt N, Vahedi-Faridi A, Tajkhorshid E & Boron WF (2021). Carbon dioxide transport across membranes. Interface Focus 11, 2.

Mitchell JK (1830). On the penetrativeness of fluids. Am J Med Sci 7, 36–67.

Mitchell JK (1833). Art. IX. On the penetration of gases. Am J Med Sci 13, 100–112.

Moll W (1968). The influence of hemoglobin diffusion on oxygen uptake and release by red cells. Respir Physiol 6, 1–15.

Moss FJ, Zhao P, Salameh AI, Taki S, Wass AB, Jacobberger JW, Huffman DE, Meyerson HJ, Occhipinti R & Boron WF (2025). Role of channels in the O₂ permeability of murine red blood cells II. Morphological and proteomic studies. *bioRxiv*; DOI: 10.1101/2025.03.05.639962.

Nakhoul NL, Davis BA, Romero MF & Boron WF (1998). Effect of expressing the water channel aquaporin-1 on the CO₂ permeability of *Xenopus* oocytes. Am J Physiol 274, C543–548.

Nicolson P & Roughton FJW (1951). A theoretical study of the influence of diffusion and chemical reaction velocity on the rate of exchange of carbon monoxide and oxygen between the red blood corpuscle and the surrounding fluid. Proc R Soc Lond, B, Biol Sci 138, 241–264.

Overton E (1895). Über die osmotischen Eigenschaften der Zelle in ihrer Bedeutung für die Toxikologie und Pharmacologie. Z Phys Chem 22, 189–209.

Overton E (1899). Ueber die allgemeinen osmotischen Eigenschaften der Zelle, ihre vermutlichen Ursachen und ihre Bedeutung für die Physiologie. Vierteljahrsschr Naturforsch Ges Zürich 44, 88–135.

Ponder E (1948). Hemolysis and Related Phenomena. New York: Grune & Stratton. Available at: http://hdl.handle.net/2027/wu.89047892849.

Popel AS (1989). Theory of oxygen transport to tissue. Crit Rev Biomed Eng 17, 257–321.

Richardson SL, Hulikova A, Proven M, Hipkiss R, Akanni M, Roy NBA & Swietach P (2020). Single-cell O₂ exchange imaging shows that cytoplasmic diffusion is a dominant barrier to efficient gas transport in red blood cells. PNAS 117, 10067–10078.

Roughton FJ, Adair GS, Barcroft J, Goldschmidt G, Herkel W, Hill RM, Keys AB & Ray GB (1936). The thermochemistry of the oxygen-haemoglobin reaction: Comparison of the heat as measured directly on purified haemoglobin with that calculated indirectly by the Van’t Hoff Isochore. Biochem J 30, 2117–2133.

Roughton FJW (1932). Diffusion and chemical reaction velocity as joint factors in determining the rate of uptake of oxygen and carbon monoxide by the red blood corpuscle. Proc R Soc Lond B 111, 1–36.

Roughton FJW (1964). Transport of oxygen and carbon dioxide. In Handbook of Physiology, pp. 767–825. American Physiological Society.

Somersalo E, Occhipinti R, Boron WF & Calvetti D (2012). A reaction-diffusion model of CO_2_ influx into an oocyte. J Theor Biol 309, 185–203.

Stannett V (1978). The transport of gases in synthetic polymeric membranes—an historic perspective. J Membrane Sci 3, 97–115.

Strutwolf J, Zhang J, Barker AL & Unwin PR (2001). Effect of phospholipids on the kinetics of dioxygen transfer across a 1,2-dichloroethane/water interface. Phys Chem Chem Phys 3, 5553–5558.

Vandegriff KD & Olson JS (1984*a*). A quantitative description in three dimensions of oxygen uptake by human red blood cells. Biophys J 45, 825–835.

Vandegriff KD & Olson JS (1984*b*). The kinetics of O_2_ release by human red blood cells in the presence of external sodium dithionite. J Biol Chem 259, 12609–12618.

Waisbren SJ, Geibel JP, Modlin IM & Boron WF (1994). Unusual permeability properties of gastric gland cells. Nature 368, 332–335.

Williams JB & Kutchai H (1986). Use of a membrane-bound fluorophore to characterize diffusion boundary layers around human erythrocytes. Biophys J 49, 453–458.

Wroblewski S (1879). On the nature of the absorption of gases. Nature 21, 190–192.

Yamaguchi K, Nguyen-Phu D, Scheid P & Piiper J (1985). Kinetics of O2 uptake and release by human erythrocytes studied by a stopped-flow technique. J Appl Physiol (1985) 58, 1215– 1224.

Yap EW & Hellums JD (1987). Use of Adair Four-Step Kinetics in Mathematical Simulation of Oxygen Transport in the Microcirculation. In Oxygen Transport to Tissue IX, Advances in Experimental Medicine and Biology, pp. 193–207. Springer, Boston, MA. Available at: https://link.springer.com/chapter/10.1007/978-1-4684-7433-6_22 [Accessed May 18, 2018].

Zhao P, Moss FJ, Occhipinti R, Geyer RR, Huffman DE, Meyerson HJ & Boron W F (2025). Role of channels in the O₂ permeability of murine red blood cells. I. Stopped-flow and hematological studies. *bioRxiv*; DOI: 10.1101/2025.03.05.639948.

